# Fragment libraries designed to be functionally diverse recover protein binding information more efficiently than standard structurally diverse libraries

**DOI:** 10.1101/2022.03.18.484642

**Authors:** Anna Carbery, Rachael Skyner, Frank von Delft, Charlotte M. Deane

## Abstract

Current fragment-based drug design relies on the efficient exploration of chemical space though the use of structurally diverse libraries of small fragments. However, structurally dissimilar compounds can exploit the same interactions on a target, and thus be functionally similar. Using 3D structures of many fragments bound to multiple targets, we examined if there exists a better strategy for selecting fragments for screening libraries. We show that structurally diverse fragments can be described as functionally redundant, often making the same interactions. Ranking fragments by the number of novel interactions they made, we show that functionally diverse selections of fragments substantially increase the amount of information recovered for unseen targets compared to other methods of selection. Using these results, we design small functionally efficient libraries that are able to give significantly more information about new protein targets than similarly sized structurally diverse libraries. By covering more functional space (rather than chemical space), more diverse sets of drug leads can be generated, increasing the chances of fragment screens resulting in viable drug candidates.

## Introduction

Fragment-based drug design (FBDD) is now well-established as a powerful approach to early-stage drug discovery, and has led to success for targets that proved otherwise intractable [1]. The first stage entails screening, where libraries of fragments, compounds around a third of the size of typical drug-like molecules, are screened for binding to a protein of interest. The concept is that the small size of the fragments allows a more efficient search of chemical space and recovers more protein binding information than in traditional high-throughput screens, allowing the size of the library to be much smaller. This substantially reduces the number of experiments that need to be conducted within a screen.

Ideally, a fragment screen obtains information about the molecules or functional groups that bind the protein of interest, and the interactions they make [2]. Fragments are subsequently elaborated or combined to create larger lead molecules that can be developed into potential drugs. The better the site of interest is explored by fragments, the more insights we have into the key interactions critical for binding. As these key interactions are usually conserved upon generation of larger lead molecules [3], this translates to improved chances of a viable drug candidate being discovered.

### Design of fragment libraries

Maximising the useful information that can be extracted from each fragment screening experiment is key, and so design of fragment libraries is a major element of FBDD research [4][5][6][7][8]. There are two major aspects to library design: definition of the desired region of chemical space, and the sampling of that region of chemical space. Table S1 describes the design strategies and size of several major fragment libraries.

#### Definition of desired chemical space

Fragment libraries tend to be built from molecules that adhere to the ‘rule of three’ [5]: fragments that have a molecular weight below 300Da, fewer than three hydrogen-bond donors and acceptors, fewer than three rotatable bonds and a cLogP of three or below. Heavy atom counts tend to be limited to below 20 [2]. These rules aim to limit the structural complexity of the fragments so that only one or two efficient interactions with the protein target are required for binding. Compounds containing toxicophores or highly reactive groups are not included [9], as these would not be appropriate fragments to use in the development of drug leads. Fragment libraries also tend to prioritise chemical tractability [10] and/or the availability of analogues [11] to enable fast and easy follow-up experiments. Such fragments that are ideal for subsequent lead development have recently been referred to as ‘social fragments’ [12], and are contrasted with ‘unsocial fragments’ which have limited or no synthetic pathways for elaboration, and no analogues.

Historical experimental results can be incorporated to further guide their definition of desired chemical space. The SpotXplorer library [13] was designed to contain pharmacophores that have been observed to commonly bind protein hotspots [14], based on a comprehensive analysis of the PDB [15]. A library designed recently [16] used past experimental results to develop a machine learning model that generated novel fragments that contain the characteristics of fragments that bind multiple targets, referred to as ‘privileged fragments’.

Libraries may also have a particular intent, for example to target certain protein classes or to fulfil specific properties. Examples of such properties include high Fsp3 character [17], 3D shape [18], the ability to form covalent bonds with the target [19], protein-protein interface binding character [20] or natural product resemblance [17].

#### Sampling of desired chemical space

To avoid the synthetic challenges presented by the design of novel compounds, most libraries are made using previously available fragments, whether available commercially or in-house. In these cases, a catalogue of fragments that lie within the desired chemical space is generated, and fragments are selected from this in a way that maximises the structural or shape diversity of the library. A common approach is to use molecular fingerprints such as ECFP [21], MACCS [22] or USRCAT [23]. A fingerprint is generated for each fragment, and a maximin-derived algorithm [24] (such as the RDKit MaxMin picker) is used to select the most structure- or shape-diverse fragments. For example, the DSiP library [25] (the successor to [10]) uses USRCAT fingerprints while the F2X libraries [11] uses MACCS fingerprints to maximise structural and shape diversity.

Another method used to achieve structural diversity is to cluster fragments based on structure or functional groups. Representatives of each cluster can then be selected for the final library. An advantage of this approach is that clusters that cover more attractive chemical space can be sampled more often than those that are less desirable. The 3D shape diverse library [26] is an example of a library employing this strategy.

There are a few libraries which consist of novel fragments that were designed and synthesised specifically for the library [27][7]. The most common aim of such libraries is to address the historic uneven coverage of chemical space. Final compounds for the library are generally selected to ensure synthetic feasibility, low similarity to commercially available fragments and to maximise shape diversity of the final library [28].

In nearly all cases, chemists are reported to be the final gatekeeper of selection, using visual inspection [16][29][30]. This indicates that algorithmic approaches are never trusted to completely select the fragments, and makes it difficult to learn from these final decisions, which are rarely fully documented or quantified.

#### Influence of HTS library design on fragment library design

To assess the effectiveness of current library design methods, it is useful to first understand how these strategies came about. The emphasis on structural diversity on library design appears to have its origin in the design of HTS libraries [31], where the large chemical space made computable metrics imperative in the selection of compounds. Experimental approaches like diversity-oriented synthesis [32] also built up on this principle. The premises were transferred to fragment library design, but to our knowledge, the underlying assumptions were never rigorously interrogated.

Gordon *et al*. [31] proposed that HTS screening libraries should be iteratively redesigned based on the results of screening campaigns. This has been applied to several libraries to guide their definition of ‘attractive chemical space’, however only knowledge of which fragments produced hits in past experiments was utilised, with 3D information from the protein-fragment structures being ignored. For example, an analysis of results from screens using the Astex 2012 fragment library [33] showed a need to focus on fragments with 10-14 heavy atoms and ensure that larger fragments are not overly complex. The Vernalis library was analysed using the results of 12 fragment screening campaigns [29] and found that compounds with slightly lower molecular weight had higher hit rates. AstraZeneca observed the high rate of project failure between 2002 and 2008 [34] and used the results to iteratively improve the library in several ways: fragments that were prone to decomposition, highly reactive or deemed ‘unattractive’ for follow-up chemistry by medicinal chemists were all removed, while pharmacophoric and structural diversity analyses were employed to ‘fill in’ gaps in chemical space.

### Functional activity of fragment libraries

Using only binary hit or miss results does not tell us whether these frequently hitting fragments are giving us diverse information about a target, so may fail in achieving the primary aim of a fragment screen of thoroughly exploring the binding site of a protein of interest. It is known that the molecular structure of a fragment does not accurately predict the interactions it is able to make with a protein [35]: similar fragments may have diverse functional activity (e.g. they bind to different protein environments) while structurally diverse fragments may have very similar activity [36]. Thus, the number of hits cannot be used as a proxy for quantity of information.

Most library designs aim to elucidate as much information as possible about a protein’s binding site(s). However, so far it has been difficult to establish which or indeed whether any fragment libraries achieve this. Now that crystallographic fragment screens are routine, we can use primary data of protein-fragment interactions to assess whether structurally diverse libraries are behaving in a functionally diverse manner.

To establish whether fragment libraries designed to be structurally diverse have functional redundancy, we examined a set of 11 diverse targets that have all been screened against the majority of fragments in the DSiP library by XChem [37]. The same set of fragments have been tested on the same targets, generating full data of what bound and how, as well as which fragments didn’t bind. Using this data we describe an approach for analysing functional redundancy within a fragment library, and examine the relationship between structurally diverse and functionally diverse fragment libraries. Our findings suggest that structurally diverse fragment libraries do not necessarily exhibit any more functional diversity than randomly selected libraries. On the other hand, by selecting functionally diverse fragment libraries, we show that the information recovered for unseen targets is substantially improved compared with using randomly selected or structurally diverse fragments.

## Materials and Methods

In this section we describe how we rank fragments by their ability to give us the most information about key interactions, and how we test the capacity of fragments we ranked highly to recover information about unseen targets.

### The XChem Dataset

Structures of 524 protein-fragment complexes were used, from 11 targets and 347 unique fragments. Each target had between 10 and 80 fragment-bound structures. The targets were extremely diverse, with a maximum global pairwise sequence identity of 8%. No two targets shared a CATH class [38]. Table S3 contains descriptions of the seven openly available targets, and the fragment-bound structures of these are available on the Fragalysis platform [39]. Full lists of fragments screened on the these targets are included within the Supplementary information.

To ensure that our analyses were not biased by fragments only observed a few times, we selected those fragments which had been tested on at least 9 of our 11 targets. This resulted in 717 fragments being included, of which 370 had not bound to any targets. The remaining 347 fragments had bound to between 1 and 4 targets. We selected fragments that had been test on at least 9 targets as a balance between coverage (9 out of 11) and dataset size. Requiring fragments to have been tested on 10 or more of our targets would have led to the inclusion of only 429 fragments in the dataset, of which only 235 had bound one or more targets. The fragments used in our analysis are included within the Supplementary information.

### Selection of functionally diverse fragments

#### Definition of functional activity

We define the functional activity of a fragment as the interactions it makes with the protein. This definition is used to prioritise our understanding of a fragment’s ability to make interactions rather than the structure of the fragment itself. The interactions in each structure within the dataset were calculated using ODDT’s InteractionFingerprint module [40], which generates a binary fingerprint for each protein-fragment structure. This method, referred to in this study as ‘residue IFP’, calculates up to eight types of interaction between the fragment and each residue in the protein (see Table S2 for details of interactions types). We also adapted the InteractionFingerprint module to output the interactions between the fragment and each atom of the protein, resulting in what we will refer to as the ‘atomic IFP’ method.

#### Ranking of fragments based on novelty of functional activity

The ranking protocol aims to identify fragments that add the most information about interactions a target can make. We start with a library size of one and append fragments one by one, each time including the fragment that adds the largest amount of novel information to the current library. As several fragments may add identical amounts of information, we repeat this 100 times, randomly shuffling the order of the fragment list before each run. At each library size, we calculate the number of interactions recovered compared to all the interactions from the full screen. The mean fractional recovery rate at each library size is calculated, along with the standard deviation. The code used to rank fragments is available at https://github.com/oxpig/fragment-ranking.

### Other methods of fragment ordering for comparison

For comparison, we used two other methods of fragment ordering, including only fragments that had bound one or more targets, to match the set of fragments that were possible to rank. The first of these was a random control, where we shuffled the fragments into a random order, repeating this 100 times and calculating the mean and standard deviation of the recovery rate at each library size. We also generated a structurally diverse control, using the technique employed in the selection of the F2X-Entry library [11]: we used RDKit [22] to calculate the MACCS key for each fragment, and MaxMin picker to select a structurally diverse set of fragments for every library size tested. This was also repeated 100 times with the mean recovery rate and standard deviation taken.

### Testing fragment ranking protocol on unseen targets

To test the effectiveness of ranking fragments by their functional information, we tested the protocol on each target in the dataset, using a leave one out test.

For each target, we ignored the results of its own screen, ranked the fragments using the other targets, and calculated the recovery of information on this previously ignored target. This was run 100 times, and the mean amount of information recovered was calculated, along with the standard deviation. These values were also calculated for randomly ordered and structurally diverse fragment libraries.

Only fragments that had bound to previously seen targets could be ranked, so once this library size was exceeded, ‘dummy fragments’ were used, which did not recover any new information from the protein. This resulted in 100 sets of fragment libraries at every size from 1 to 225 for each of the library types. For each method, the mean information recovered was calculated, along with the standard deviation of this information recovery.

## Results

Structural data from fragment screens of 11 unrelated protein targets bound to 717 fragments were used in this study. Protein-ligand interaction fingerprints (IFPs) were calculated for each structure, between fragment atoms and protein residues (residue IFP) and between fragment atoms and protein atoms (atomic IFP). For both types of IFP the fragments were ranked based on the novel interactions they made with all or a subsection of protein targets (see Methods). These rankings were used to group fragments for analysis, defined in Table 1.

**Table 1:**
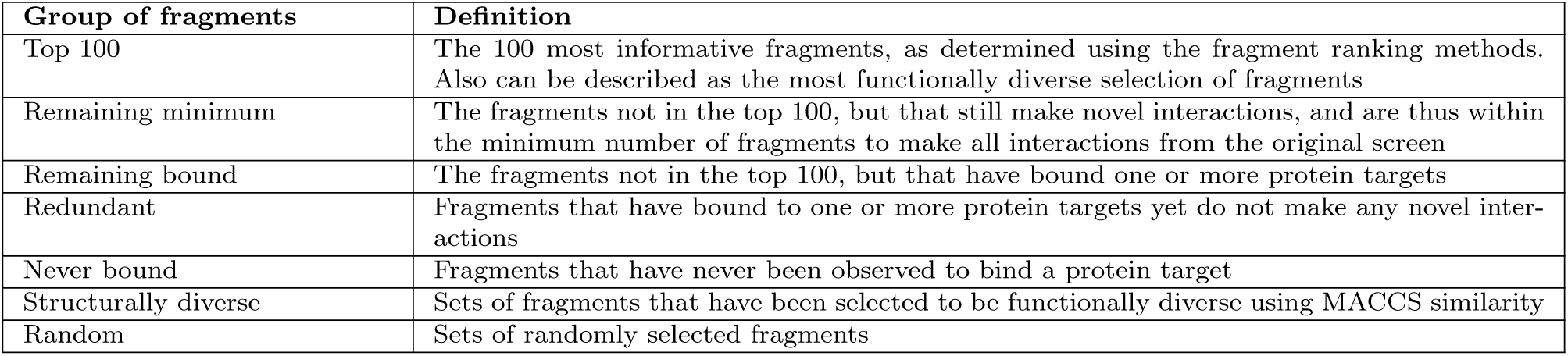
Definitions for the various groups of fragments used in this analysis.

### Structurally diverse fragments can make overlapping interactions

We compared the molecular similarity (calculated using ECFP2 fingerprints [21]) to the residue IFP (our measure of functional activity) similarity for fragments bound to 11 highly diverse protein targets (Table S3). Fragments that are structurally very different (ECFP2 similarities as low as 0.02) bound to the same location on a target and make one or more of the same interactions, leading to an IFP similarity above zero (figure 1a). Across the set we found 90 pairs of structurally distinct fragments that formed identical interactions. An example of this structural dissimilarity but functional similarity is shown in figure 1b, where three distinct fragments form identical interactions with the protein. The molecular fingerprint (ECFP2) similarity of these three fragments ranges between 0.27 and 0.34, which would be considered appropriately diverse for inclusion in conventional libraries.

**Figure 1:**
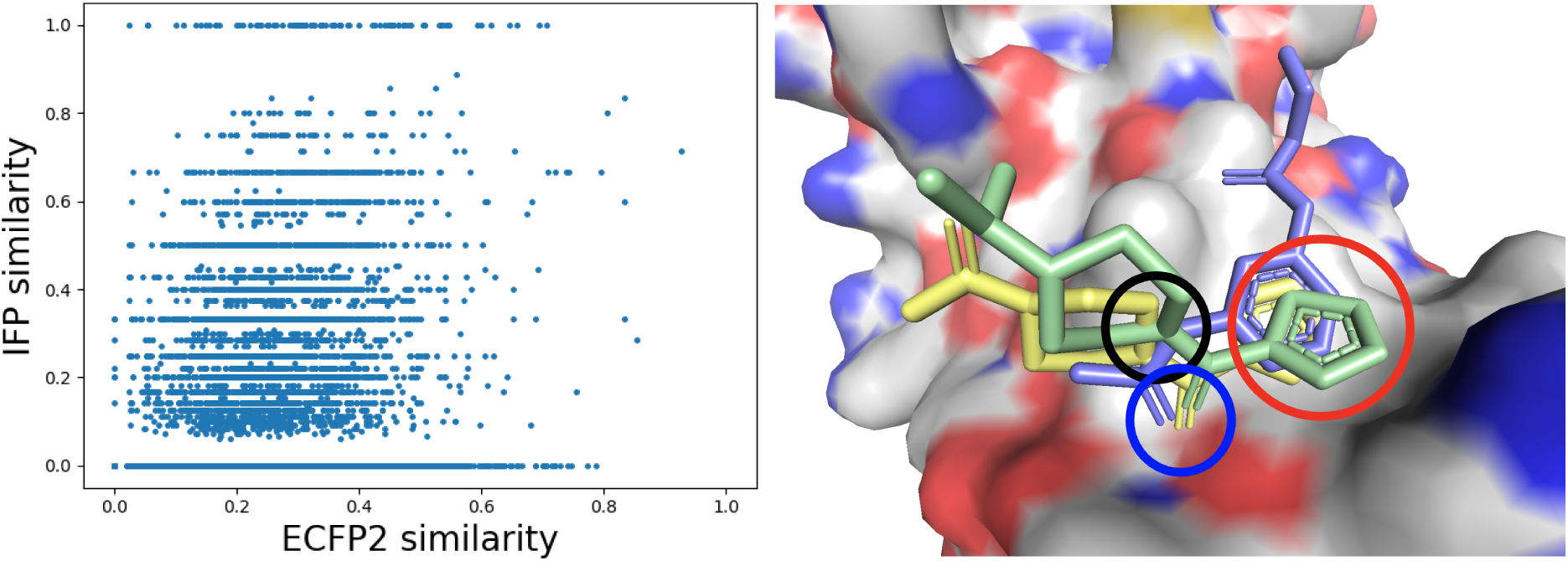
Very diverse fragments make identical interactions. **(a):** Molecular (ECFP2) similarity compared to functional (IFP) similarity. Each point represents a pair of fragments that bind to the same target. There is no direct correlation between ECFP2 (molecular) similarity and IFP (functional) similarity, showing that structural diversity of fragments is not predictive of functional diversity. **(b):** An example of three fragments (pale blue, green and yellow) which make the same interactions with target TBXTA (grey). The atoms circled in bright blue (carbonyl oxygens) are making hydrogen bonds with two residues; atoms circled in black (nitrogens) are making a hydrogen bond with a single residue; atom groups circled in red are making a hydrophobic interaction with a single residue.

While the 90 pairs of fragments that form identical interactions only account for 0.27% of all fragment pairs, over a fifth (21.7%) of fragment pairs shared at least one common interaction. To explore the possibility that fewer fragments could make the same interactions with the targets, we assessed which fragments made the most novel interactions, and calculated the minimum number of fragments required to explore all interactions across all targets.

### Ranking of fragments reveals redundancy in interactions

Fragments were ranked by the number of novel interactions (residue-level and atomic-level) that they made with the 11 targets in the dataset (Table S3). Two libraries, one randomly ordered and another structurally diverse, were generated for comparison to the functionally diverse libraries (see Methods for details of how these were prepared). Each of these three methods of ordering fragments was repeated 100 times, and the mean fraction of information recovered at each library size across all runs was calculated (Figure 2).

**Figure 2:**
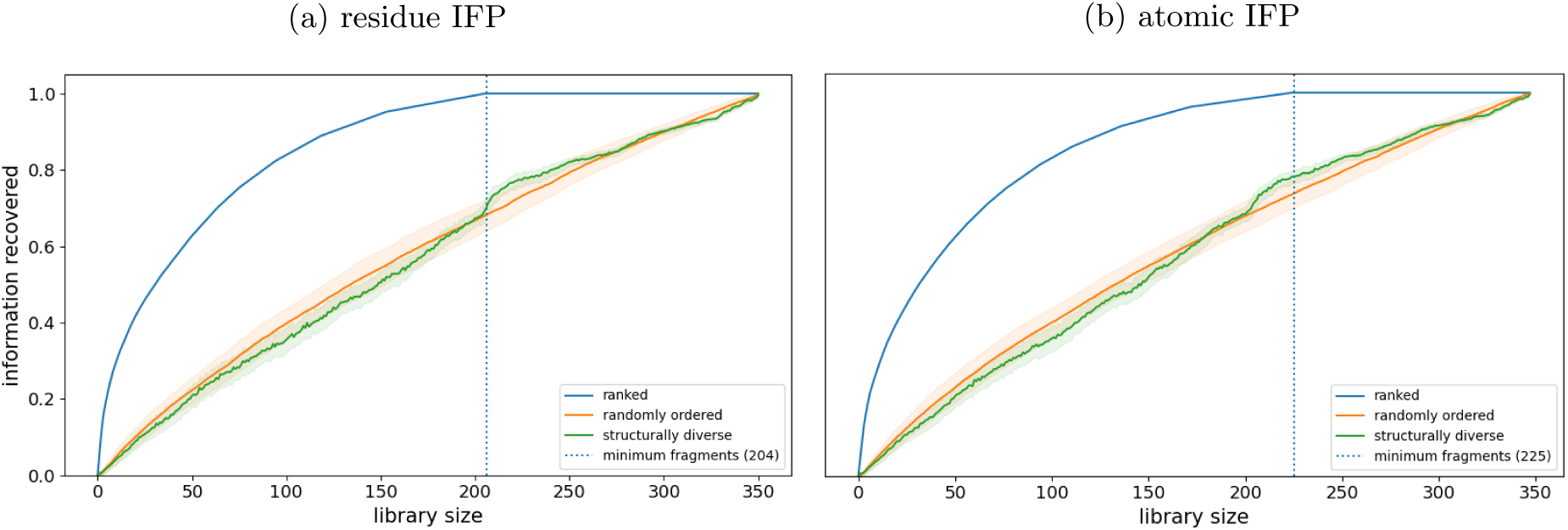
Ranked fragment libraries recover information at smaller library sizes than other methods. Blue dotted line indicates minimum number of fragments required to recover all information. **(a):** fragments have been ranked using the residue IFP method. **(b):** fragments have been ranked using the atomic IFP method.

Interactions were recovered at smaller library sizes for the functionally ranked library compared with either the random or the structurally diverse library. On a residue level, all interactions with targets were recovered using only the 204 top-ranked fragments of the 347 that were bound to at least one target (Figure 2a); for atomic level of interactions, 224 fragments were required (Figure 2b). This result shows that by selecting the fragments in this way, a 224-fragment screen could recover the same information as a randomly selected 347-fragment screen. It was expected that the ranked libraries (using both residue and atomic IFP) would recover information at smaller library sizes due to the method of ranking, however it is notable that structurally diverse libraries do not recover information at library sizes any smaller than the randomly selected libraries. The lack of difference between structurally diverse and random libraries may be because the fragments used are already relatively structurally diverse.

To assess whether different libraries were selecting similar fragments, the mean number of fragments in common between libraries of 100 fragments was calculated. The functionally diverse and the structurally diverse libraries had 26 fragments in common, similar to the 25 fragments in common between the functionally diverse and randomly selected fragments. This indicates that the level of similarity between the functionally diverse and structurally diverse libraries is little more than random. Conversely, the functionally diverse libraries generated using the residue IFP and atomic IFP methods have 86 fragments in common.

### Functionally diverse compounds exhibit different chemical properties to non-binding fragments

To explore the relationship between particular chemical properties and a fragment’s rank, we split the DSiP fragment library into three sets: the top-ranked 100 fragments (those that are most informative), the 247 remaining bound fragments (fragments that had bound to one or more proteins), and the 370 fragments that had never bound to a target (Figure 3). This was performed on sets generated by both residue and atomic IFP methods, and as a comparison, chemical properties were also calculated for a structurally diverse set of 100 fragments.

**Figure 3:**
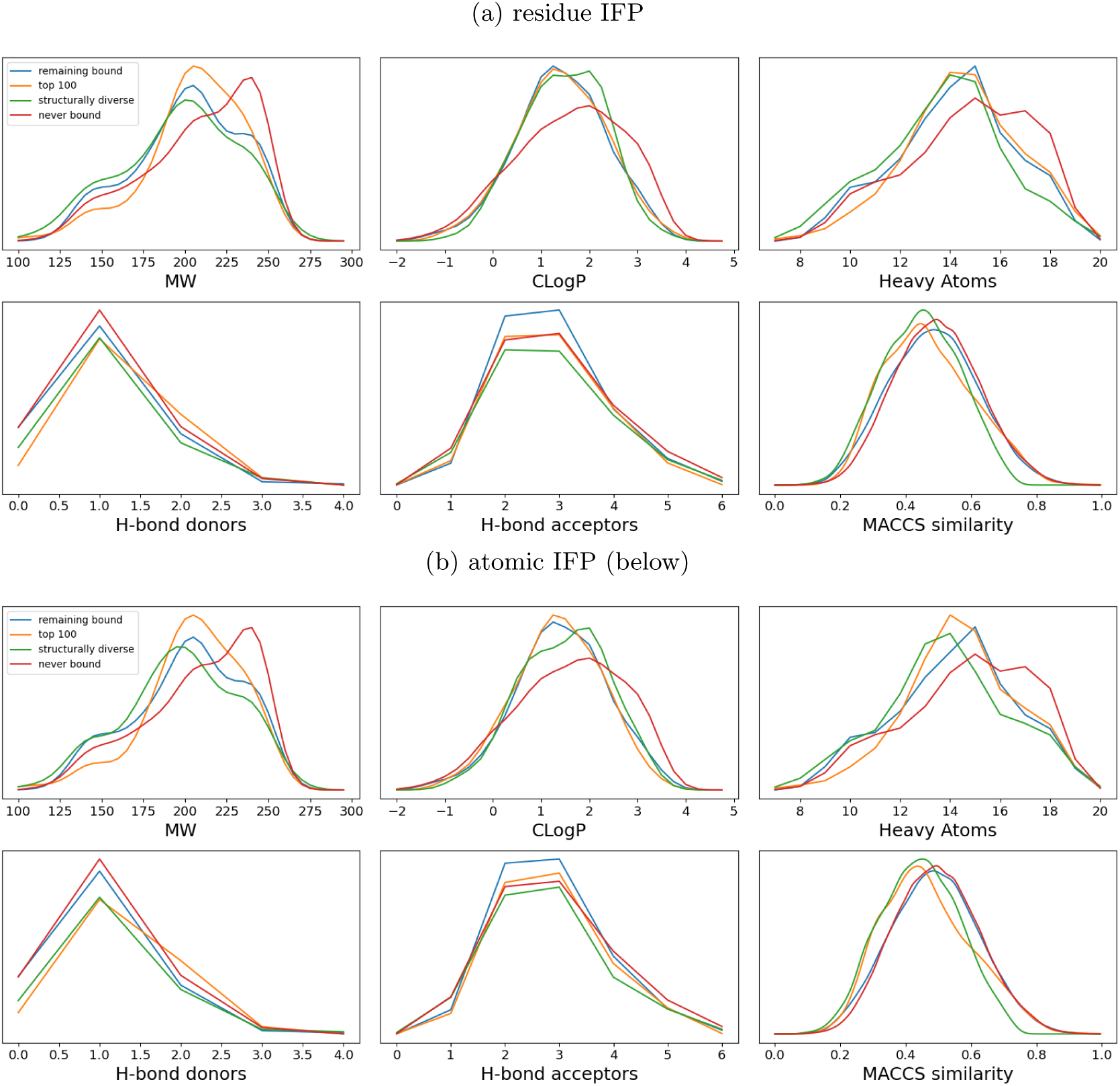
Fragments that have never bound to a target (red) are more likely to have higher MW and heavy atom counts, along with lower solubility. Fragments have been categorised into the ‘top 100’ and ‘remaining bound’ groups based on our functional ranking (see Methods). Fragments that bound no targets make up the ‘never bound’ set, while a set of structurally diverse fragments are shown for comparison. Various properties of these sets of fragments are compared. **(a):** fragments have been ranked using the residue IFP method. **(b):** fragments have been ranked using the atomic IFP method. **(a) and (b):** Medium-sized fragments are most likely to be highly informative.

Fragments ranked in the top 100 functionally diverse were more likely to have a molecular weight (MW) around 200 for both residue and atomic IFP methods, while the ‘other bound’ groups of fragments were slightly more likely to have a MW under 175. For all other properties, there was no substantial difference between groups of fragments that had bound one or more targets. Fragments that had not been observed to bind any target were more likely to have a higher MW (over 225), a higher heavy atom count (17 or above), and a slightly lower solubility than those that had bound one or more targets. As expected, the fragments selected as a diverse subset had lower MACCS similarity with each other compared with the overall library and the highly-ranked fragments.

### Promiscuous binders are not necessarily the most informative fragments

To examine whether a fragment that bound multiple targets (a promiscuous fragment) was likely to be selected as a highly informative fragment, we compared the number of hits (targets bound) for three sets of fragments: the 100 top-ranked fragments, the remaining minimum fragments (fragments that would be required to recover all information from the original screen), and the fragments that could be removed without losing any information (redundant fragments). While the top-ranked sets were made up mostly of fragments that had bound multiple targets, some fragments that bound to 3-4 targets were excluded in favour of those that had only bound to a single target (Figure 4).

**Figure 4:**
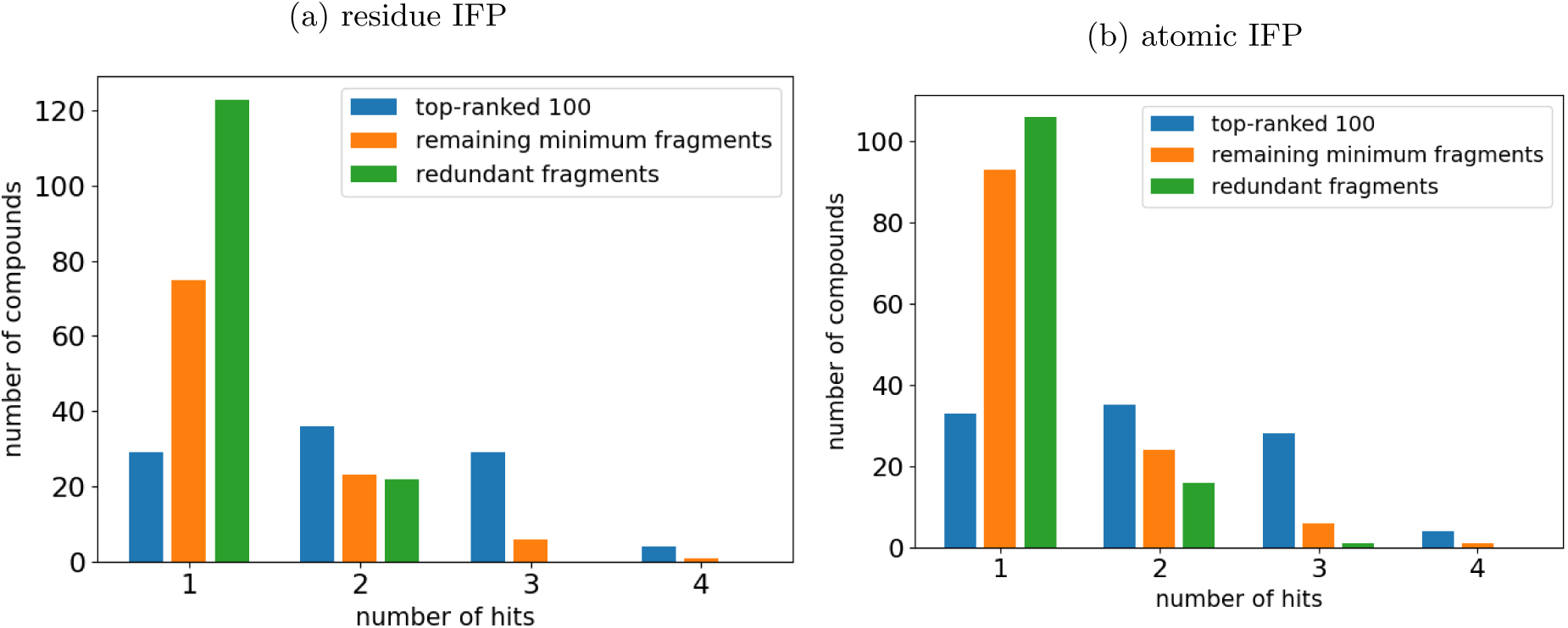
Number of targets bound by fragments in the DSiP library, grouped by their position when ranked. The top-ranked 100 fragments are compared to those remaining fragments that give us novel interactions, and to fragments that make only redundant interactions. Promiscuous fragments (e.g. 3 or 4 hits) are not always selected as the most informative, whereas some fragments with only 1 hit are ranked in the top 100. **(a):** fragments ranked using the ‘residue IFP’ method. **(b):** fragments ranked using the ‘atomic IFP’ method.

### Functionally diverse fragments recover information more efficiently from unseen targets

The results above show that a functionally diverse fragment set contains different fragments to a structurally diverse one. We investigated whether using such functionally diverse fragments is an effective strategy to more efficiently get information about unseen targets. To do this, we performed a leave one out test (see Methods for more details).

To compare the information recovery for each target when using functionally diverse fragments with the other methods of fragment selection, we analysed the information recovered from each target at a library size of 100 fragments (Figure 6). We also calculated the the fractional improvement when using the functionally diverse fragments compared with random and structurally diverse fragments (Figure S1).

We compared the methods of fragment selection across all unseen targets. The mean values of information recovery across all targets are shown in Figure 5b. On average, the functional information about the unseen target was recovered more efficiently using the functionally diverse fragments than the random libraries.

**Figure 5:**
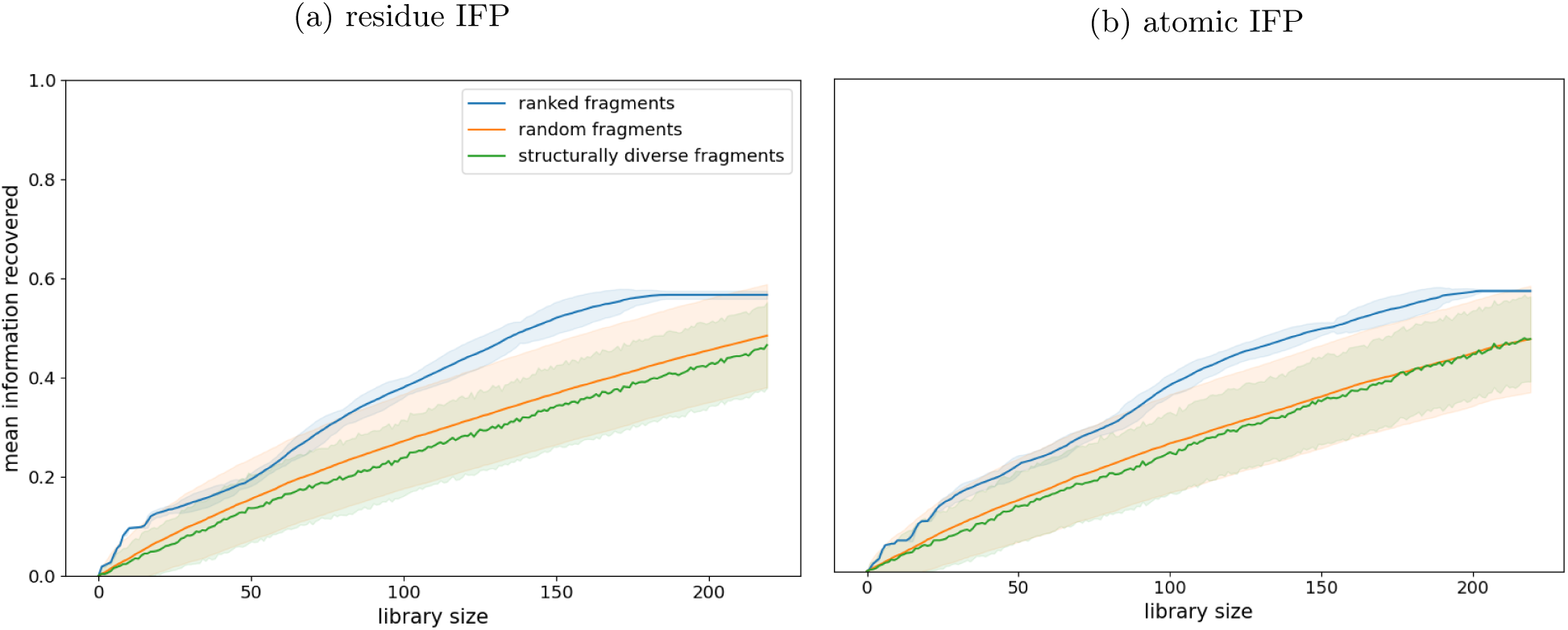
Ranked fragment libraries show superior information recovery for unseen targets at every library size. The recovery of information for each target when unseen was calculated 100 times. The mean across each run for each target was calculated, and the mean of these values was taken. This value is shown at each library size, with error clouds showing 1 standard deviation across 100 runs. **(a):** fragments have been ranked using the ‘residue IFP’ method. **(b):** fragments have been ranked using the ‘atomic IFP’ method.

**Figure 6:**
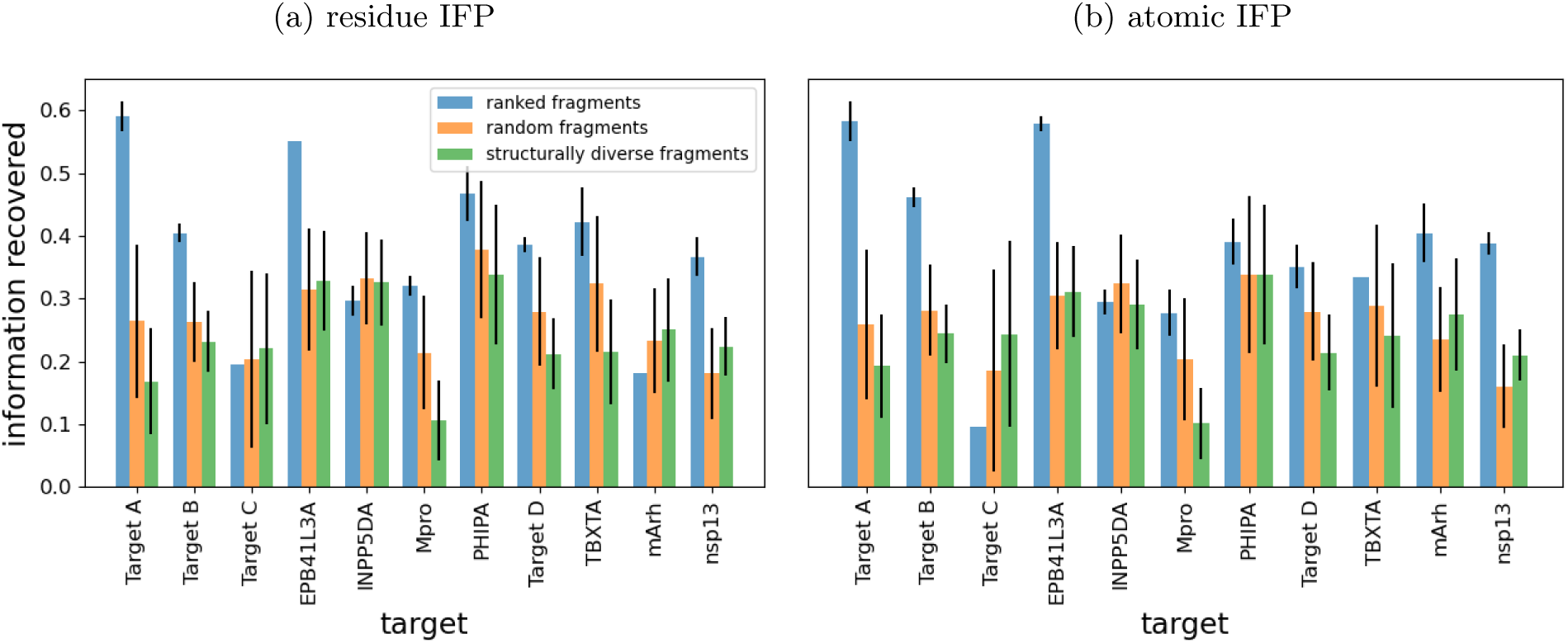
Top-ranked fragments show superior information recovery compared with random fragments and structurally diverse fragments. For each target, average information recovery when using the top-ranked 100 fragments over 100 runs is shown. Error bars show 1 standard deviation. Lack of error bars show no variability in the result. **(a):** fragments have been ranked using the ‘residue IFP’ method. **(b):** fragments have been ranked using the ‘atomic IFP’ method.

### Targets contribute differently to prediction of important fragments for unseen targets

The targets in this dataset are very diverse (the maximum pairwise global sequence identity is 8%), but some fragments do make similar interactions with different targets. To assess the impact of each target on the effectiveness of a fragment set for giving information about an unseen target, we removed the previously seen targets one by one, ranked only on the remaining 9 targets, and took the top-ranking 100 fragments as the functionally diverse library. The recovery of interactions was compared with the original (when results of all 10 targets were used to rank), and the factor that each target impacts the recovery of interactions for every other target was calculated. These impact scores are a measure of the similarity of the most informative fragments between two targets (figure 7a-b). Between the residue IFP method and the atomic IFP method, the scores are consistent in terms of overall effect, but the magnitudes differ. Each target positively impacts some targets while negatively impacting others. Particular targets seem to be negatively affected by the presence of many targets at a residue level but not at an atomic level (for example, targets Target C and mArh).

**Figure 7:**
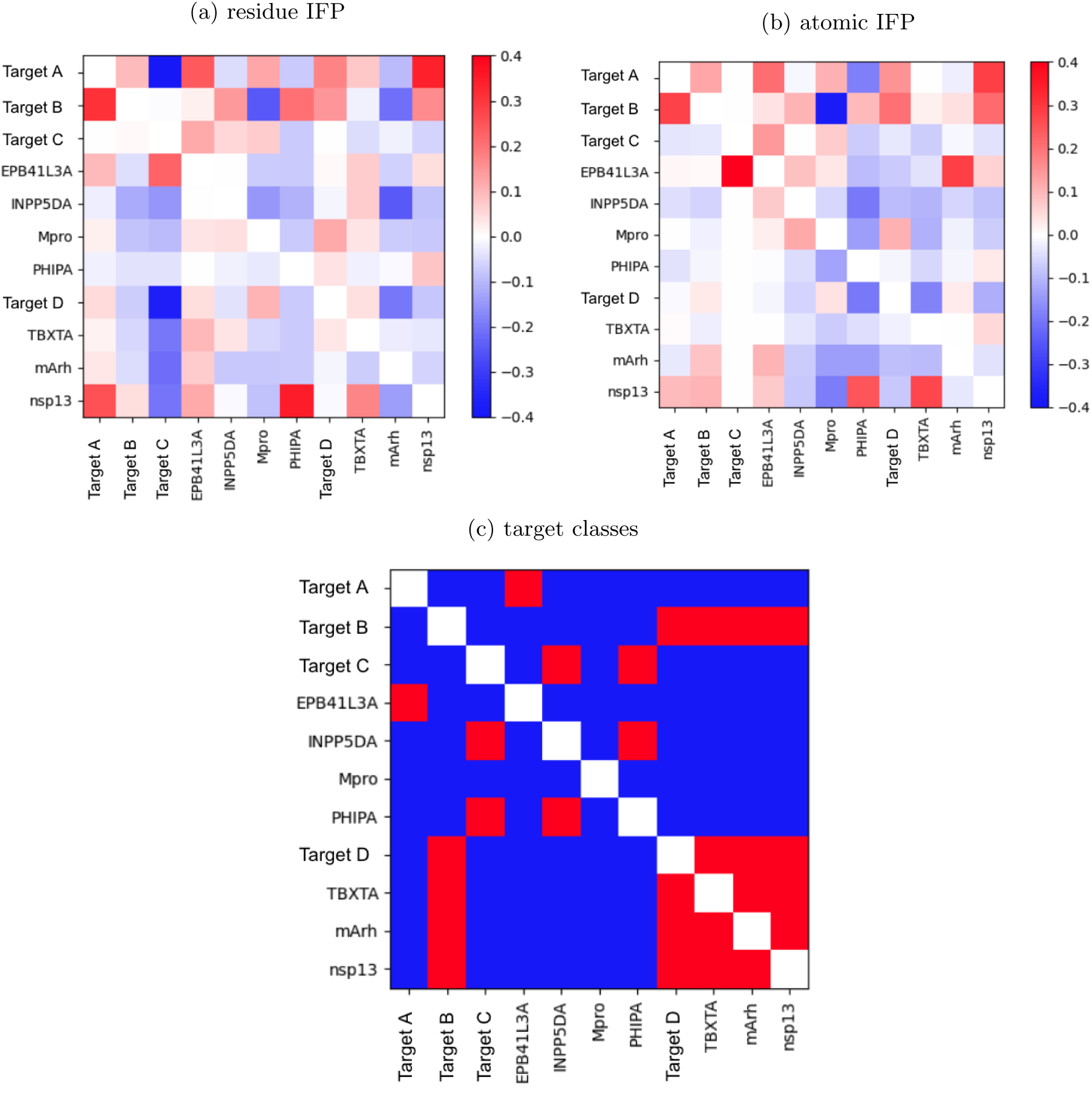
The impact of each target on every other target’s information recovery at library size of 100 fragments. For example, including nsp13 improves PHIPA’s recovery of information by a factor of 0.2-0.4 compared with only using the remaining 9 targets to rank the fragments. **(a):** fragments have been ranked using the ‘residue IFP’ method. **(b):** fragments have been ranked using the ‘atomic IFP’ method. **(c):** A heatmap showing whether two targets fall into the same class. Red indicates that two targets are in the same class and blue indicates a different class.

As a high impact score is indicative of similar fragments making interactions with the targets, we assessed which targets were in the same classes (phosphatase/kinase, protease, nucleic or other). This is shown in Figure 7c, however it does not appear that there is a correlation between these two factors. This confirms that there are no two targets with similar fragment binding activity, and yet by ranking fragments on previously seen targets, we can improve information recovery on unseen unrelated targets.

## Discussion

In agreement with previous work [36], we found that structurally diverse fragments are able to make similar or even identical interactions. Additionally, we have shown that by defining fragments as the interactions they make with all targets, fragment libraries can be selected based on their functional diversity. Such libraries are able to make more diverse interactions with previously unseen targets, and thus improve the information recovered by an average of 50% and a maximum of 250% across all targets tested, compared with traditionally designed fragment libraries.

These findings suggest that selection based on structural diversity is not the optimal strategy when diverse functional information is desired. Given that suitable experimental data is now available for many fragments bound to multiple targets, it is feasible to explore new approaches to select fragments to screen previously unseen targets.

### Ranking fragments shows redundancy in interactions

In order to select the fragments that show the most diverse functional activity, we ranked fragments by the number of novel interactions they made with 11 targets. This showed that some fragments make far more novel interactions than others. We analysed the most informative fragments (those that were most highly ranked) and compared them to less informative fragments and those that have never been seen to bind. Our analysis of the most informative fragments is broadly in agreement with previous work [41] and suggests that fragments with molecular weights between 175 and 225 (and heavy atom counts between 12 and 16) perform optimally in fragment screens, perhaps as they can make multiple interactions without being structurally too complex. Additionally, while promiscuous binders (fragments that bind multiple targets) are more likely to be considered as highly informative fragments, their promiscuity alone is not enough to guarantee this. This supports our hypothesis that using only information on which fragments have a high hit rate is not the most effective strategy for library redesign.

### Functionally diverse fragments recover information more efficiently from unseen targets

We then proceeded to study the potential of the above described ranking protocol as a novel method for selecting fragments to screen on unseen targets. We tested whether a set of fragments that exhibited functional diversity in previously screened targets were more efficient in information recovery than a random set of fragments or a structurally diverse set of fragments on an unseen target. Both residue IFP and atomic IFP ranking methods achieved better information recovery than comparison methods: the residue IFP method improved recovery by a maximum of 250% and an average of 50%, and the atomic IFP method improved recovery by a maximum of 210% and an average of 50%. This result indicates that the functional redundancy of the DSiP fragment library remains across diverse targets and that designing the library in a functionally diverse way can lead to more information being generated for a target from a screen compared with traditionally designed libraries.

The atomic IFP method gives slightly more consistent results than the residue IFP method. This suggests that the particular atom within a residue that a fragment interacts with is important, as the atomic IFP method captures this information whereas the residue IFP method does not.

### Different targets contribute differently to prediction of important fragments for unseen targets

Finally, we set out to understand whether particular targets contributed disproportionately to the performance of our method, thus indicating that there were targets with similarity in their fragment binding activity within the dataset. By testing the impact of each target’s screening results on the recovery of information for every other target in a 100-fragment screen, we scored the similarity between the most informative fragments between each pair of targets. Each target positively impacts some targets while negatively impacting other. Some targets are mostly negatively impacted by others at a residue level while being unaffected or positively affected at an atomic level. As a measure of binding similarity, we compared this to substrate class of protein, however there was little to no correlation. Considering the complexities of protein-ligand binding behaviour, it is unlikely that a simple predictor of this behaviour exists, and so further research would be required to explore potential ways to predict such similarities between targets. Additionally, as we would expect the impact of each target on each other target will change as more results are included and artefacts due to the relatively small dataset used here are reduced.

## Conclusions

Currently, the most common strategy for fragment library design is to select the most structurally diverse set of fragments from those that lie in the desired chemical space, without considering structural results from previous fragment screens. In this study we have shown that libraries designed on the basis of functional diversity recover information more efficiently from unseen targets than traditionally designed structurally diverse libraries.

Even with a limited dataset, we have proven the potential for functionally diverse fragment selections to substantially improve the information recovered from fragment screens. With a much larger dataset, it would be possible to use our methods to select functionally diverse fragments that can reliably and significantly outperform traditional library selection methods. This ability to better explore the interactions in protein binding sites would allow a larger number of diverse lead compounds can be developed, thus improving the chances of a fragment screening campaign producing a viable drug candidate.

## Supporting information

Fragments included in analysis

Full lists of compounds screened

## Acknowledgements

A.C. is supported by the Engineering and Physical Sciences Research Council (EPSRC) (Reference: EP/N509711/1) and Diamond Light Source. Crystallographic data were collected by the XChem group and collaborators at Diamond Light Source on beamline I04-1. In addition to the XChem group, we extend thanks to the following groups for allowing us to include their data in this analysis: Evgenii M. Osipov, Ali H. Munawar, Steven Beelen and Sergei V. Strelkov (KU Leuven, Belgium); Martin Noble and Natalie Tatum (Newcastle University, United Kingdom); Norbert Sträter, Susanne Moschütz, Konstantin Richter and Renato Weiße (Universität Leipzig, Germany); Xiangrong Chen, Mark S. Roe, Laurence H. Pearl and Antony W. Oliver (University of Sussex, United Kingdom).

## Supplementary Information

**Table S1:** Many of the fragment libraries currently in use. Some are used privately, while others are commercially available. Empty cells are indicative that the method has not been published.

| Library Name | Chemical space | Sampling strategies | Size | Reference |
| --- | --- | --- | --- | --- |
| DSiP | poised | shape diversity (USRCAT) | 768 | [25] |
| F2X | pharmacophore | ph4 diversity (MACCS) | 96 | [11] |
| SpotXplorer | hotspot exploration | ph4 + interaction clustering | 96 | [13] |
| Fraglites | halogenated fragments with paired H-binding motifs |  | 25 | [42] |
| PepLites | halogenated peptidomimetics |  | 25 | [43] |
| Cambridge 3D | 3D and poised | diversity oriented synthesis | 137 | [44] |
| York 3D | 3D, substituted aliphatic heterocycle | shape diversity | 106 | [28] |
| Leeds 3D | 3D, natural product-like scaffolds, high sp<sup>3</sup> |  |  | [45][17] |
| MiniFrag | Astex’s pharmacophores, ultra-low-MW | ‘chemical diversity’ | 81 | [46] |
| Astex | general | chemical and structural diversity | 2000 | [33] |
| CovHetFrag | Small heterocyclic electrophiles | chemical properties | 141 | [47][48] |
| CysElectrophile | Cysteine covalent library |  | 993 | [19] |
| Novartis | general | structural diversity, clustering | 1408 | [49] |
| AZ | general xrc | structural diversity (ECFI) | 384 | [50] |
| Chembridge | large set | subsets: solubility, fluorine, bromine, structurally diverse | 15000 | [51] |
| Asinex | ”BioDesign” - synthetically feasible, NP-like and common NP ph4 |  | 20061 | [52] |
| Life Chemicals | known bioactives | diversity, Fsp3, target family-specific | 8000 | [53] |
| Prestwick | Fragments of approved drugs |  | 1456 | [54] |
| Selcia |  |  | 1214 | [55] |
| Timetec |  | structural diversity |  | [56] |
| Zenobia | general inc. PPI | shape diverse | 352 | [57] |
| Bilsland | ML-generated privileged fragments | FeCo, novelty, ‘examination by medicinal chemists’ | 741 | [16] |
| Essential | initial screen, frequently reported hits |  | 320 | [26] |
| High fidelity | MedChem tractable | diversity selection using clustering algorithm | 1920 | [26] |
| Fluorinated | <sup>19</sup>F NMR |  | 1000 | [26] |
| Covalent | Diverse covalent |  | 6210 | [26] |
| Fully-functionalized | Photoaffinity labeled fragments |  | 2000 | [26] |
| NP-like | NP-like | scaffold frequency analysis | 4160 | [26] |
| 3D shape-diverse | 3D diverse | K-mean clustering, centroids taken | 1200 | [26] |
| PPI | peptidomimetics |  | 3600 | [26] |
| Single ph4 | ro3-like | variety of ph4s and scaffolds | 1500 | [26] |
| Carboxylic acid | specific protein targets |  | 4000 | [26] |
| Halogen-enriched | halogen bonding motifs | diversity | 3000 | [26] |
| Fully-functionalized probe | Efficient exploration of novel protein targets w/ photoaffinity probes |  | 640 | [26][58] |
| EU openscreen | availability of parent molecules within larger collection |  | 1056 | [59] |
| Maybridge | pharmacophorically-rich | clustering and centroids selected | 2500 | [60] |
| Vernalis | general | diversity of ph4 triangles | 1063 | [29] |
| Pfizer | general | diversity and visual inspection | 2592 | [30] |

**Table S2:** The eight types of protein-ligand interaction detected by ODDT’s InteractionFingerprint module [40].

| Bit | Interaction type |
| --- | --- |
| 0 | hydrophobic contacts |
| 1 | aromatic face to face |
| 2 | aromatic edge to face |
| 3 | hydrogen bond (protein as hydrogen bond donor) |
| 4 | hydrogen bond (protein as hydrogen bond acceptor) |
| 5 | salt bridges (protein positively charged) |
| 6 | salt bridges (protein negatively charged) |
| 7 | salt bridges (ionic bond with metal ion) |

**Table S3:**
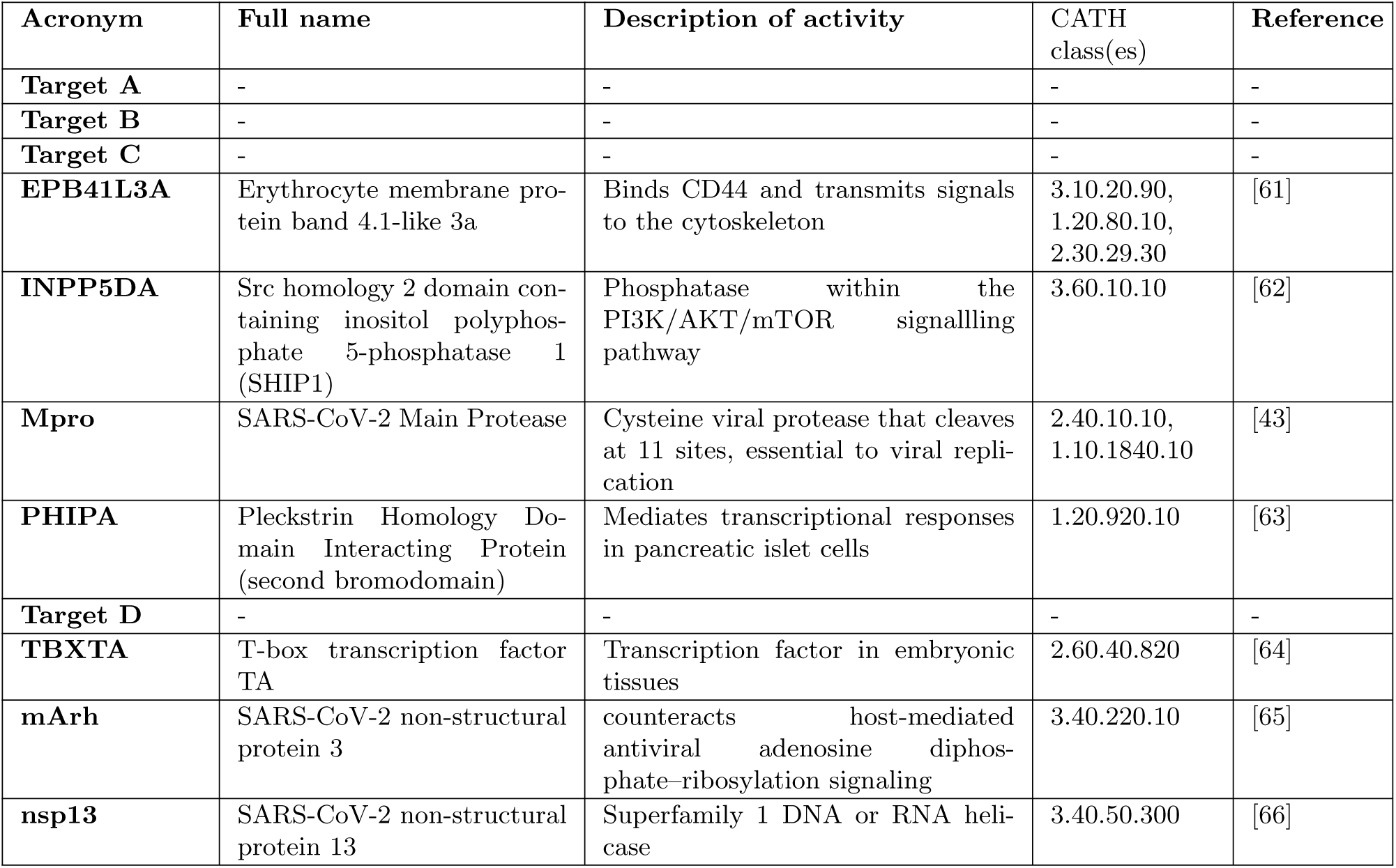
Descriptions of each target used in this study. CATH classes obtained through sequence-based searching of CATH [38] at https://www.cathdb.info/.

| Acronym | Full name | Description of activity | CATH class(es) | Reference |
| --- | --- | --- | --- | --- |
| Target A | - | - | - | - |
| Target B | - | - | - | - |
| Target C | - | - | - | - |
| EPB41L3A | Erythrocyte membrane protein band 4.1-like 3a | Binds CD44 and transmits signals to the cytoskeleton | 3.10.20.90, 1.20.80.10, 2.30.29.30 | [61] |
| INPP5DA | Src homology 2 domain containing inositol polyphosphate 5-phosphatase 1 (SHIP1) | Phosphatase within the PI3K/AKT/mTOR signalling pathway | 3.60.10.10 | [62] |
| Mpro | SARS-CoV-2 Main Protease | Cysteine viral protease that cleaves at 11 sites, essential to viral replication | 2.40.10.10, 1.10.1840.10 | [43] |
| PHIPA | Pleckstrin Homology Domain Interacting Protein (second bromodomain) | Mediates transcriptional responses in pancreatic islet cells | 1.20.920.10 | [63] |
| Target D | - | - | - | - |
| TBXTA | T-box transcription factor TA | Transcription factor in embryonic tissues | 2.60.40.820 | [64] |
| mArh | SARS-CoV-2 non-structural protein 3 | counteracts host-mediated antiviral adenosine diphosphate-ribosylation signaling | 3.40.220.10 | [65] |
| nsp13 | SARS-CoV-2 non-structural protein 13 | Superfamily 1 DNA or RNA helicase | 3.40.50.300 | [66] |

**Figure S1:**
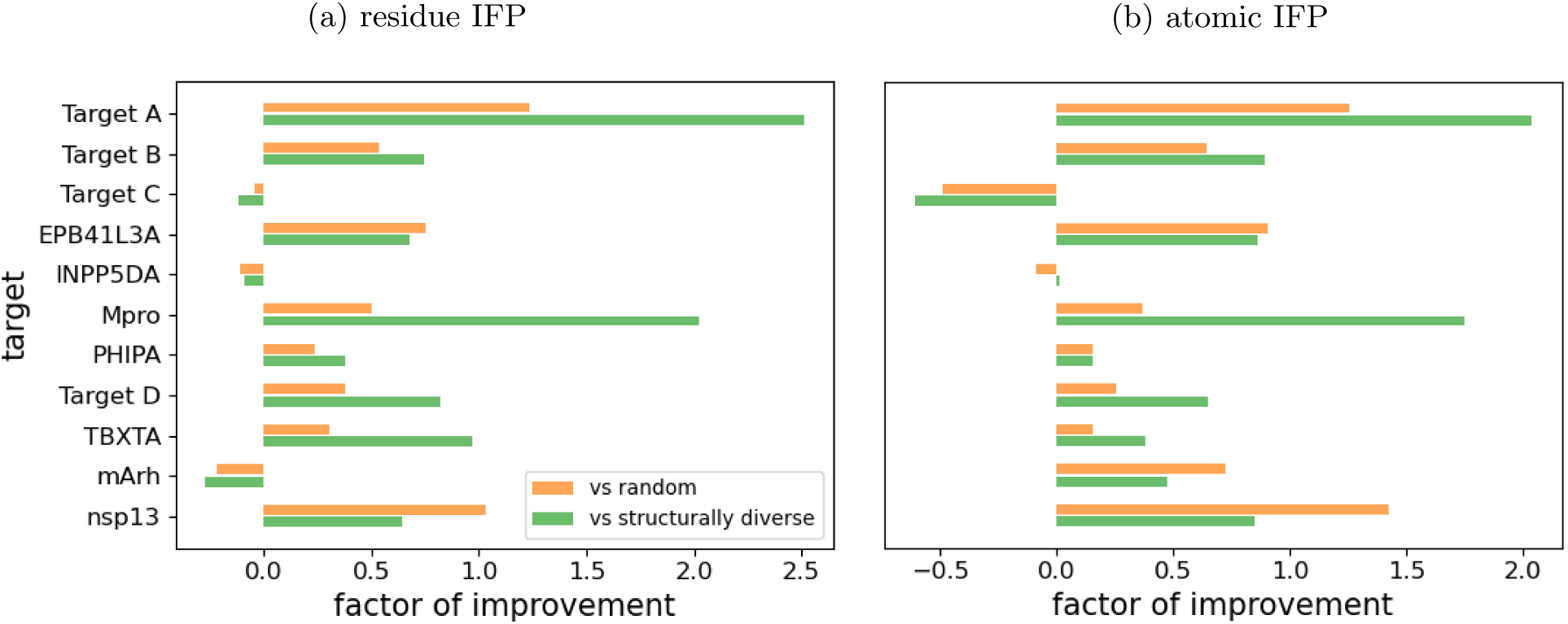
Factor of information improvement when using 100 functionally diverse fragments compared with random and structurally diverse libraries. **(a):** fragments have been ranked using the residue IFP method. **(b):** fragments have been ranked using the atomic IFP method.

