## Supplementary material for "Fragment libraries designed to be functionally diverse recover protein binding information more efficiently than standard structurally diverse libraries": Fragments included in analysis

| SMILES | | | |
|---|---|---|---|
| CC(C)CC(=O)NCc1ccc2c(c1)OCO2 | Oc1cccc(CN2CCSCC2)c1 | CCNc1ccc(C)nn1 | CCOC(=O)c1sc(N(C)C(C)=O)nc1C |
| Cc1cc(CN(C)C(=O)NC2CC2)no1 | Cc1nccc(N2CCC(N)CC2)n1 | CCCC(=O)NCc1nc2ccccc2[nH]1 | CC(CO)(CO)NC(=O)Nc1ccccc1 |
| CN(C)C(=O)C1(C(=O)O)CC1 | CCCC(=O)Nc1cccnc1Cl | CNc1nccn1 | CC(=O)Nc1ccc(CN2CCOCC2)cc1 |
| C1CC(Cc2nc(C3CC3)no2)CCN1 | OCc1cn(-c2ccc(Cl)cc2)nn1 | OCCN1CCOCC1 | CN(CC(=O)O)C(=O)c1ccccn1 |
| CC(=O)N1CC(C)Oc2c(F)cccc21 | CC(CO)N(C)c1ncc(Cl)cn1 | CNCC(=O)Nc1cc(C)ccn1 | Cn1cc(Oc2ncncc2Cl)cn1 |
| CC(O)C(=O)NCc1ccccn1 | CNC(=O)C1CCCc2sc(C)nc21 | Cc1ccc(CN2CCS(=O)(=O)CC2)cc1 | COc1cc(CN)cc(OC)c1OC |
| CC(C(N)=O)N1CCN(S(C)(=O)=O)CC1 | COc1ccc(CNC2CCCC2)cc1 | CCc1ccc(NC(=O)c2cncn2)cc1 | CC(C)(N)CN1CCCC1 |
| Cc1cc(C)nc(NC(=O)Nc2ccccc2)c1 | Oc1ccccc1-c1nc(-c2ccccc2)no1 | Cn1nnc(NC(=O)c2ccc(Cl)cc2)n1 | CS(=O)(=O)NCCc1ccc(F)cc1 |
| CCNC(=O)CN1CC(C)OCC1C | CC1CCOC1C(=O)Nc1cnns1 | Cc1ccc(OCC(=O)Nc2cc(C)on2)cc1 | CC(=O)NCC1CNCCO1 |
| Cc1nc(C(=O)N2CCCC2)c(C)s1 | Cn1nccc1C(=O)NCc1cccs1 | COC(=O)COc1ccc(C#N)cc1 | O=C(COc1ccccc1)N1CCCC1 |
| CC(C)Nc1nc2ccccc2[nH]1 | COCC(=O)Nc1c(C)cc(C)cc1C | NC(=O)c1cnc(C(F)(F)F)nc1 | CC(C)NC(=O)CN1CCCNCC1 |
| Cc1ccc(CNC(=O)C2CCC2)cn1 | CN1CCN(C(=O)CO)CC1 | CC(=O)Nc1ccc(CC(N)=O)cc1 | CC1(C)CN(C(=O)NCC2CC2)C1(C)C |
| CC(=O)NC1CNc2ccccc2C1 | NC(=O)c1ccc(OCc2ccccc2)cc1 | CCc1nnc(NC(=O)c2cncn2)s1 | CCNc1ncccc1C |
| COc1cccc2c1CCCC(=O)N2 | CCC(=O)NCc1nnc2n1CCCC2 | CNC1CCCCC1S(C)(=O)=O | OC1CCN(Cc2ccsc2)CC1 |
| CCNC(=O)N1CCN(C(C)=O)CC1 | Cc1nnc(N2CCCCC2)s1 | CNS(=O)(=O)c1cccc(C)c1F | c1ccc(CN2CCC(n3cccc3)CC2)cc1 |
| CCOC(=O)N1CCOCC1 | CC1COCCN1Cc1ccccc1F | COc1cccnc1C#N | O=C(NCCc1ccc(O)cc1)c1ccccn1 |
| Cc1ccc(N)c(=O)n1C | CCNCc1cccc2[nH]ccc12 | COC(=O)CNC(=O)c1cc(C)on1 | O=C(Nc1ccc2[nH]c(=O)[nH]c2c1)c1ccco1 |
| O=C(NCCc1ccncc1)NC1CCCCC1 | Cc1ccc(-c2noc(CNC(C)C)n2)cc1 | CC(=O)N1CCCC(C(N)=O)C1 | CCc1ccccc1NC(=O)c1snnc1C |
| COC(=O)c1cccc(NS(C)(=O)=O)c1 | CC(=O)c1ccc(NC(=O)CN2CCCC2)cc1 | COc1ccc(CNC(=O)c2csnn2)cc1 | Cc1n[nH]c(C)c1-c1ccccc1N |
| CC(=O)c1cccc(NC(=O)C(F)(F)F)c1 | Cc1nnc(NC(=O)N2CCOCC2)s1 | N#CCC(=O)NC1CCCCC1 | CNC(=O)Nc1nc2ccc(OC)cc2s1 |
| CNS(=O)(=O)Cc1cc(C)on1 | CN1CCN(c2ccc(C#N)cc2)CC1 | CCOC(=O)CN1CCCCC1 | CN(C)S(=O)(=O)c1cn[nH]c1 |
| C[C@H](O)CNCc1c(Cl)cccc1Cl | O=C(NCCC1=CCCCC1)c1cnccn1 | Cc1c(Cl)cccc1NC(=O)[C@@H]1CCCO1 | O=C(Nc1nccs1)[C@@H]1C[C@H]1c1ccccc1 |
| Cc1cc(C(=O)NC(C)C)on1 | O=C(NCC1CC1)Nc1ccc2c(c1)CCO2 | COC(=O)Nc1cc(C)n(-c2ccccc2)n1 | Cc1occc1C(=O)Nc1cc(C(C)(C)C)on1 |
| Cc1ncc(CN2CCNCC2)s1 | COc1ccccc1-c1nnco1 | CC(N)C(=O)Nc1cc(C(F)(F)F)n[nH]1 | CC(=O)N1CCCC(O)(C(=O)O)C1 |
| CCNCc1cn(C)nn1 | CC1CCNC(C(N)=O)C1 | O=C(CN1CCCC1)Nc1cccc(Cl)c1 | c1cc(-c2ccc3c(c2)OCO3)n[nH]1 |
| CC(Oc1ccc(Cl)cc1)C(N)=O | CCc1ccc(OCC(=O)N2CCCCC2)cc1 | CC(O)CCn1cccn1 | Cc1occc1C(=O)N1CCc2ccccc21 |
| Cc1nnc(CN2CCC=C(F)C2)s1 | CC1CCN(C(=O)c2cc3ccccc3o2)CC1 | CCOC(=O)CN1CCS(=O)(=O)CC1 | COc1ccc(Cl)cc1C(=O)NCC(C)C |
| CNCc1ccc(C(F)(F)F)cc1 | c1cc(CN2CCNCC2)no1 | O=C(O)c1cccc(NC(=O)c2ccno2)c1 | CC(Oc1ccc(-c2ccccc2)cc1)C(=O)O |
| Cc1ccc(C(=O)Nc2nnc[nH]2)cc1C | CCC(=O)Nc1c(C(N)=O)oc2ccccc12 | NC(=O)c1ccc(NC(=O)[C@@H]2CCCO2)cc1 | COc1ccccc1N1CCN(CC#N)CC1 |
| Fc1ccc(CNC[C@H]2CCCO2)cc1 | CCc1cc(=O)n2nc(NC)sc2n1 | Cc1noc(NC(=O)C2CCCC2)n1 | CC(=O)N(C)c1cccnc1 |
| Nc1ccccc1N1CCOCC1 | Cc1ccc(OCC(=O)N2CCOCC2)c(C)c1 | C[C@H](O)CNC(=O)Nc1ccccc1 | c1csc(CNCc2ccncc2)c1 |
| C[C@@H](CO)Nc1ncc(Cl)cc1F | CCOc1cccc(C(=O)Nc2nnn(C)n2)c1 | O=C(c1ccco1)N1CCN(C(=O)C2CC2)CC1 | CNC(=O)CN(C)C1CCCCCC1 |
| CC(=O)N1CCN(CCCc2ccccc2)CC1 | Cn1cc(C(N)=O)c(C(F)F)n1 | CC(=O)Nc1ccc(F)c(C(=O)O)c1 | COC(=O)[C@@H]1C[C@@H](O)CN1C(=O)c1ccco1 |
| CNC(=O)c1ccccc1NC(=O)c1ccco1 | COC(C)c1nc(-c2cnoc2C)cs1 | C1CCC(NCC2CCCO2)C1 | CCOC(=O)c1cn[nH]c1 |
| COc1ccc(CN)cc1OC | CCC(=O)Nc1nc(C)cs1 | COc1ccc(Nc2ncnc3[nH]cnc23)cc1 | CC(C)C(N)C(=O)N1CCOCC1 |
| CCOc1ccccc1N1CCNCC1 | COc1ccc(CNC(=O)CC#N)cc1 | C1CN(CC2CCOCC2)CCN1 | CCS(=O)(=O)Nc1ccccc1F |
| Cc1[nH]cnc1CN1CCc2ccccc2C1 | N#CCC(=O)Nc1ccc(Cl)cc1 | NC(=O)C1CCN(C(=O)Nc2ccccc2)CC1 | COC(=O)c1sccc1NC(=O)C1CC1 |
| Cc1ccc(CNC(=O)Cc2cccs2)cc1 | CNCc1nnc2ccccn12 | NC(=O)N1CCN(c2ccc(F)cc2)CC1 | Cc1cccc(C(=O)N(C)C)c1C |
| CN(C)Cc1ccc(C#N)cc1 | CC(C)(O)CN1CC[C@H](N)C1 | CC(=O)NC(C)(C(=O)O)C1CC1 | CC[C@H](CO)NC(=O)Nc1ccc(Cl)cc1 |
| CC(=O)Nc1ncc(Cc2cccc(C)c2)s1 | CC(=O)N1CCC(N)CC1 | CN(C)C(=O)OCCn1ccnc1 | C[C@H](O)CNCc1ccc2c(c1)OCO2 |
| O=C(CCCc1ccccc1)N1CCOCC1 | CC(N)c1cn(-c2ccccc2)nn1 | Cc1ccccc1CC(N)=O | c1ccc2sc(N3CCOCC3)nc2c1 |
| CC(C)COc1ccc(C(=O)N2CCCC2)cc1 | CC(=O)Nc1ccc(Oc2ncccn2)cc1 | CC(C)c1ncc(Cl)c(C(N)=O)n1 | CS(=O)(=O)c1nccn1Cc1cscn1 |
| CC(=O)N1CCC(C#N)CC1 | O=C(Cc1ccccc1)N1CCc2ccccc21 | CCOc1ccc(C(=O)Nc2cc(C)on2)cc1 | CCn1cc(CCO)cn1 |
| c1nc(N2CCC2)c2[nH]cnc2n1 | COc1ccc(C(=O)NCC(N)=O)cc1 | CC(=O)Nc1cccc(F)c1C(=O)O | Nc1ccccc1OCc1ccccc1 |
| Cn1cc(CNC(=O)c2ccco2)cn1 | Nc1nccc(-c2ccc(Cl)cc2)n1 | Cc1cccc(NC(=O)Nc2nc(C)cs2)c1 | Cc1nc(CN2C[C@@H](F)C[C@H]2CN)cs1 |
| NC(=O)Nc1ccc(-c2ccccc2)cc1 | CCCn1nccc1-c1ccncc1 | Cc1cc(C)nc(Nc2ccccc2)n1 | CC(=O)Nc1cc(C)n[nH]1 |
| CC(O)CNC1CCCCC1 | Cc1cccc(N2CCNC(C)C2)n1 | CC1CCCN(C(=O)CO)C1 | CC(=O)NC(C(=O)O)C1CCOCC1 |
| COCC(=O)Nc1nc(-c2ccccc2)cs1 | CS(=O)(=O)NC1(C(=O)O)CCCCC1 | CS(=O)(=O)CC(O)c1ccccc1 | NC(=O)CN1CCN(C(=O)c2ccco2)CC1 |
| c1cc(NCC2CCCO2)ccn1 | CC(N)c1ccc(N2CCOCC2)cc1 | CCS(=O)(=O)N1CCC(C(=O)O)CC1 | Cc1ccc(C(N)=O)c(F)c1 |
| CCNC(=O)Nc1cc(C)on1 | O=C(NCc1ccco1)NC1CCCCC1 | O=C1CN(C(=O)COc2ccccc2)CCN1 | CC1COCCN1CCN |
| CCOc1ccc(C(N)=O)cn1 | COc1ccc2sc(N)nc2c1 | O=C(Cc1ccc(F)cc1)NC1CCCCC1 | CS(=O)(=O)NCCc1ccccc1 |
| Fc1ccccc1NCc1cc[nH]n1 | CNCC1COc2ccccc2O1 | CN1CCN(CCN)CC1C(C)C | CCN(CC)c1cc(C)nc2ncnn12 |
| CC(C)(O)CNC1CCOCC1 | CS(=O)(=O)N1CCC(O)CC1 | CC(C)(C)N1CC(O)C1 | COc1ccc(NC(=O)Nc2ccncc2)cc1 |
| O=C(Nc1ccc(O)cc1)c1ccc(Cl)cc1 | CCOC(=O)Nc1cccc(Cl)c1 | CC(=O)NCCc1ccccc1C | Cc1ccccc1OCC(=O)Nc1ccccc1 |
| COC(=O)Nc1sc(C)nc1-c1ccccc1 | CCN(CC)S(=O)(=O)c1ccc(C)cc1 | O=C(Nc1ccc(F)cc1)Nc1cccc(O)c1 | CCC(=O)Nc1cccc(N)c1C |
| Cc1cc(C(=O)Nc2ccc3[nH]ccc3c2)no1 | Cc1ccccc1OCC(=O)N(C)C | CCc1nnc(NC(=O)C(F)(F)F)s1 | Oc1ncc(NC2CCCCC2)c(O)n1 |
| Cc1nsc(N2CCOC(C)C2C)n1 | CC(=O)NCCc1c[nH]c2ccc(F)cc12 | OCC1CCN(c2ncccn2)CC1 | Cn1ccc(NCc2ccccc2)n1 |
| CNCc1cccc(OC)n1 | CS(=O)(=O)Nc1ccc(C(N)=O)cc1 | O=C(Nc1ccc(O)cc1)N1CCOCC1 | CC(C)NC(=O)N(C)C1CCS(=O)(=O)C1 |
| CCNC1CCN(c2ccncc2)CC1 | O=C(CSc1ccccc1)NC1CCCCC1 | Oc1ccc(CN2CCc3ccccc3C2)cc1 | CC(C)Nc1cccnc1 |
| Cc1cncc(NCc2ccncc2)c1 | CCN(C)C(=O)c1cccc(N)c1 | CC(C)N(C)S(=O)(=O)N1CCNC(=O)C1 | O=C(c1ccc2c(c1)OCO2)N1CCCCCC1 |
| Cc1n[nH]c(C)c1S(=O)(=O)N(C)C | CN(CCO)c1ccccn1 | CNC(=O)CCC1CCCCC1 | Cc1ccc(-c2nc(N)sc2C)cc1C |
| CS(=O)(=O)NCC1CCNCC1 | CC(=O)N1CCCNCC1 | N[C@@H]1CCCN(C(=O)CCC(F)(F)F)C1 | CC(=O)Nc1ccc(C)c2c1CCCN2 |
| CS(=O)(=O)Nc1ccc(-c2ccccc2)cc1 | COCC(=O)Nc1cccc(NC(C)=O)c1 | CNCc1ccncc1 | CCC(=O)Nc1ccc2[nH]c(=O)[nH]c2c1 |
| CC(=O)N(C)c1ccccc1C(=O)O | CC(C)(C)CC(=O)Nc1ccc(F)cc1F | CNS(=O)(=O)c1ccccc1Cl | O=C(c1c(F)cccc1F)N1CCCC1 |
| CC(=O)Nc1cccc(C(=O)O)c1C | CC(N)c1ccc(-c2ncncc2)cc1 | COc1ccc(C(=O)Nc2ccccc2F)cc1 | CCOC(=O)c1ccc(F)cc1F |
| O=C(NC1CC1)c1ccc2c(c1)OCO2 | O=C(NC[C@H]1CCCO1)N1CCOCC1 | Clc1cccc(CN2CCOCC2)c1 | CC1CN(S(=O)(=O)N(C)C(C)C)CCO1 |
| CN(C)c1ccc(CNn2cnnc2)cc1 | C1CC(CNC2CC2)CO1 | CCC(=O)Nc1ccc(C)cc1O | COCc1nnc(N)s1 |
| O=S1(=O)CCCN1Cc1cccc(F)c1 | O=C(Nc1ccccc1)Nc1cccnc1 | Cc1nc(CC(=O)N2CCOCC2)cs1 | Cc1nc(-c2ccccc2)no1 |
| CN(C)C(=O)Nc1ccc(S(N)(=O)=O)cc1 | CN(C1CCCCCC1)S(C)(=O)=O | Cc1noc(C(N)C(C)C)n1 | CC(=O)NCC1(O)CCCC1 |
| CC1CCCN(C(=O)c2cscn2)CC1 | c1cc(-c2noc(C3CCCCC3)n2)ccn1 | CNCc1cnn(-c2ccccc2)c1 | CN1CCC[C@@H]1CO |
| O=C(NCc1ccccc1)c1cccc(F)c1 | COc1cccc(C(=O)Nc2ccccc2F)c1 | CCC(=O)NC1CCNC1 | CC(NC(=O)C1CCCC1)C(=O)O |
| NNC(=O)c1ccc(OCc2ccccc2)cc1 | Cc1ccsc1CNC(C)(C)CO | CS(=O)(=O)Nc1ccc(CN)cc1 | CCCc1nnc(NC(=O)c2ccccn2)s1 |
| O=C(NO)c1ccc2ccccc2n1 | CC(=O)Nc1cc2c(cc1C)OCO2 | COc1ccc(C(N)C(=O)O)cc1 | COCC(=O)Nc1ccc(O)cc1 |
| COCC(=O)NCc1ccccc1 | CCC(=O)Nc1ccc(N2CCC(C)CC2)cc1 | Cc1noc(CN2CCC(CO)CC2)n1 | Cc1cc(NC(=O)CCC2CCCCC2)no1 |
| O=C(Nc1ccon1)c1cccc(Cl)c1 | Cc1ncsc1C(=O)N1CCNC(=O)C1 | NC(=O)c1ccccc1NC(=O)c1ccncc1 | CCN1N=C(C(F)(F)F)CC1=O |
| O=C(CC1CCCCC1)Nn1cnnc1 | O=C(NCCc1cccs1)C1CCCCC1 | NC(=S)Nc1ccccc1OC(F)(F)F | CC(=O)N1CCC(NCCc2ccccc2)CC1 |
| CNC(C)c1ccc(F)cc1 | Cc1ccsc1C(=O)O | N#Cc1ccc(Cn2cnc3ccccc32)cc1 | COc1ccc(C(N)c2ccccc2)cc1 |
| NCc1cc(-c2ccccc2)no1 | Cc1[nH][nH]c(=O)c1CCO | Cc1sc2ncnc(N(C)C)c2c1C | C[C@@H]1C[C@H]1C(=O)NCCc1ccc(Cl)cc1 |
| COc1cccc2[nH]ccc12 | Cc1nn(C)c(C)c1CN1CCNCC1 | COCC1CN(c2ccccn2)CCO1 | CN(C(=O)[C@H]1CC1(C)C)c1cccnc1 |
| CC1CCCCN1C(=O)c1cnccn1 | COC(=O)C1CCN(C(=O)NC(C)(C)C)CC1 | Cc1c(Cl)cccc1NC(=O)CN | Nc1ccc(S(=O)(=O)Nc2ccccc2)cc1 |
| Cc1ccccc1-n1cc(C(=O)O)nn1 | CC1CNCCN1CCO | CCC(=O)Nc1ccc(S(N)(=O)=O)cc1 | CN(C)C(=O)CNc1ccccc1 |
| CCNC(=O)c1ccc(NS(C)(=O)=O)cc1 | O=C(Nc1nnc(-c2ccco2)s1)C1CC1 | Cc1csc(NC(=O)C2CCOCC2)n1 | CCc1ccc(-c2nc(N)sc2C)cc1 |

| | | | |
|---|---|---|---|
| CC(C)c1nnc(NCC(C)(C)C)s1 | CS(=O)(=O)N1CCC[C@@H]1CN | CC(C)C(=O)NCC1CCCNC1 | COCC(=O)Nc1ccccc1 |
| Cn1cc(Cl)c(C(=O)NC2CCCC2)n1 | c1coc(CNc2nc3ccccc3[nH]2)c1 | CC(C)C(=O)NCc1nc2ccccc2[nH]1 | Clc1snnc1CN1CCOCC1 |
| CC(=O)c1ccc(OC(F)F)cc1 | CC(=O)NCc1ncc[nH]1 | CNC(=O)c1ccc(S(N)(=O)=O)cc1 | O=C(Nc1ccc(F)cc1)Nc1ccc(F)cc1 |
| Cc1ncc(CNc2cncc(F)c2)s1 | COC(=O)Nc1nc2c(C)cccc2s1 | COc1ccc(NC(=O)Nc2cccs2)cn1 | CCc1noc(C)c1C(=O)Nc1cc(C)on1 |
| CCC1CCC(NC[C@H]2CCCO2)CC1 | COc1ccc(NCc2ccccc2O)cc1 | CC(=O)c1ccc(N2CCC(C(N)=O)CC2)cc1 | CNC(C)c1cnn(C)c1 |
| N#CCN1CCN(c2cccc(Cl)c2)CC1 | O=C(NCCc1ccccc1)NC1CCCCC1 | CNC(C)c1sc(C)nc1C | CNc1nccnc1C#N |
| CC(Cc1ccc(Br)cc1)C(N)=O | CCc1csc(-c2c[nH]cn2)n1 | CC(=O)Nc1ccc(OC(=O)N2CCCC2)cc1 | N#Cc1ccc(N2CCNCC2)c(F)c1 |
| CCn1cc(C(=O)NCc2ccc(F)cc2)cn1 | CC(=O)Nc1ccncc1C(=O)O | COc1ccc(CN2CCOCC2)cc1OC | Oc1ccccc1CN1CCOCC1 |
| CC(C)Nc1ncccn1 | O=C(OCc1ccc(Cl)cc1)c1cccnc1 | COc1ccc2nc(NC(=O)C3CC3)sc2c1 | COc1ccccc1NC(=O)c1ccc(C)s1 |
| COc1ccc(NC(=O)CN)cc1 | CNC1(C(N)=O)CCCC1 | Cn1nc(C(N)=O)c2ccccc21 | CS(=O)(=O)Cc1nc2ccccc2[nH]1 |
| Cc1occc1C(=O)Nc1ccncc1 | CC(C)c1ccc(NC(=O)N2CCOCC2)cc1 | CN1CCCc2ccc(S(N)(=O)=O)cc21 | O=C(NCc1nc2ccccc2[nH]1)c1ccco1 |
| CC1CN(C(=O)c2cnsn2)C(C)CO1 | Oc1ncc(NCCc2ccccc2)c(O)n1 | CC(c1ccccc1F)N1CCNCC1 | Cc1cccc(C(=O)NCC(=O)O)c1 |
| CN1CCC[C@@H](OC(=O)c2ccco2)C1 | CC(=O)N1CCN(c2ccc(Cl)cc2)CC1 | Cc1cc(C(=O)NCC2CCCO2)ns1 | Cc1cccc(CN2CCOCC2)c1 |
| CC(C)N(C)c1ncnc2c1cnn2C | CCn1c(NC(C)=O)nc2ccccc21 | COC(=O)c1ccc(S(N)(=O)=O)cc1 | c1cnc(N2CCC3(CCOC3)C2)cn1 |
| CC1CCN(C(=O)CC(F)(F)F)CC1 | CC(C)CNC(=O)c1ccc2ccccc2c1 | CC(C)NCc1ccc(N(C)C)cc1 | COc1ccc(C)cc1NC(=O)Nn1cnnc1 |
| Cn1cc(C(N)=O)c(C2CCCNC2)n1 | Cc1cc(F)cc(S(N)(=O)=O)c1 | CCOC(=O)c1cc(-c2ccc(C)cc2)no1 | N#Cc1ccc(CNC(=O)N2CCOCC2)cc1 |
| Nc1cccc(OCc2ccccc2)c1 | Cc1nc(CN2CCC(CN)CC2)cs1 | NC(=S)Nc1ccc(OC(F)(F)F)cc1 | CC(C)(CO)NC(=O)c1cccc(Cl)c1 |
| Fc1ccccc1CNc1cn[nH]c1 | Cc1noc(C)c1C(=O)NCCC1=CCCCC1 | CC1CN(C(=O)c2cnccn2)CCO1 | OCc1ccc(-n2cnc3ccccc32)cc1 |
| O=C(C1CC1)N1CCN(c2ccc(F)cc2)CC1 | O=C(NCCc1ccc(F)cc1)c1ccco1 | NC(=O)C1CCCn2ccnc21 | O=C(O)Cc1ccc(-c2ccccc2)cc1 |
| CCN1CCN(C(=O)NC2CCCCC2)CC1 | CC(C)(C)C1CCN(C(=O)c2ccccc2)CC1 | CC(C)(CO)NC(=O)Nc1ccc(Cl)cc1 | CC(=O)Nc1ccc(NC(=O)[C@H]2CC2(C)C)cc1 |
| C(#Cc1ccccc1)CN1CCCCC1 | NC(=O)c1ccnc(NC2CC=CC2)c1 | COC(=O)N1CCN(c2ccc(F)cc2)CC1 | CNC(=O)c1c(-c2ccc(F)cc2)noc1C |
| Cc1nsc(N2CCCNCC2)n1 | Cc1nccn1Cc1cccc(C#N)c1 | CN(C)Cc1nc(C2(N)CCCC2)no1 | Cc1c(C(=O)NCC2CCC2)cnn1C |
| CC(=O)Nc1cc(C(=O)O)ccc1F | O=C(COc1ccc(F)cc1)N1CCCC1 | CCS(=O)(=O)NCc1ccc2c(c1)OCO2 | CC(C)N1CCN(C(=O)c2cccs2)CC1 |
| Cn1nnc(NC(=O)c2cc3ccccc3o2)n1 | CC(N)C(=O)N1CCCCC1C | O=C(c1c(F)cccc1F)N1CCCCCC1 | CNC(=O)CN1CCNCC1 |
| Cc1ccccc1OCC(=O)Nc1ncccn1 | CC(C)(C)NC(=O)Cn1ccc(C(F)(F)F)n1 | Cc1csc(CNC(=O)c2cc[nH]n2)n1 | CCn1cc(NC(=O)C2CCCC2)cn1 |
| CC(C)C(=O)Nc1nnn(C)n1 | CCN(CCO)c1ccc(N)cc1 | CC(NC1CC1)c1cccnc1 | CCn1cc(CNC(=O)c2cccnc2)cn1 |
| CCCn1nnnc1NC(=O)c1ccco1 | O=C(Nc1ccc2ncccc2c1)c1ccccc1 | Cc1cc(C(=O)NCc2cccs2)no1 | Cn1nnc2c1CC(C(=O)O)CC2 |
| N[C@H]1CCN(S(=O)(=O)c2ccccc2)C1 | CCNc1ccc(C#N)cn1 | Cn1ncc(Cl)c1C(=O)N1CCCCC1 | CC(C)c1ccc(C(=O)Nc2ccccn2)cc1 |
| Cc1ccc(OCC(=O)N2CCN(C)CC2)cc1 | Oc1ccc(CNC2CCCCC2)cc1 | c1nc(CN2CCNCC2)cs1 | O=C(NCc1ccco1)Nc1ccc(F)cc1 |
| CNC1CCCN(c2ccccc2)C1=O | CN(C)c1ccc(C(=O)Nc2ccncc2)cc1 | Cc1cccc(NC(=O)[C@@H]2CCCN2)n1 | CN1CCN(C(=O)c2ccc(F)c(F)c2)CC1 |
| CN1CCN(C(=O)Nc2ccccc2)CC1 | CN(CCO)S(=O)(=O)c1ccc(N)cc1 | CCOc1ncccc1C(N)=Oa | CNC(=O)c1cccc2cc[nH]c12 |
| CC(=O)c1cccc(NC(=O)N2CCOCC2)c1 | Cc1ccc(NC(=O)c2ccccc2C)c(O)c1 | C1CN(Cc2nc(C3CC3)no2)CCO1 | O=C(COC(F)F)N1CCCC2(CC2)C1 |
| c1csc(-c2ncc[nH]2)c1 | O=C(Nc1cccc(Cl)c1)C(F)(F)F | CC(=O)N1Cc2ccc(N)cc2C1 | CS(=O)(=O)NC1CCCC1 |
| OCc1ccccc1N1CCOCC1 | COC(=O)c1ccccc1NC(=O)N1CCCC1 | CCNC(=O)c1c[nH]nn1 | CC1CCN(Cc2c[nH]c3ccccc23)CC1 |
| CC(C)n1cc(C(=O)O)nn1 | O=C(CC(F)(F)F)N1CCNCC1 | CC(C)C(=O)Nc1cccc(C#N)c1 | Cc1cc(NC(=O)CSc2ccccc2)no1 |
| CC(O)c1cnn(C2CCCC2)c1 | c1ccc(NCC2CCNC2)nc1 | Cc1cc(C(=O)Nc2cnn(C)c2)no1 | NC(=O)N1CCN(C(=O)c2ccco2)CC1 |
| CC(=O)N1C[C@H]2CC[C@H]1c1ccccc12 | CS(=O)(=O)NC1CCNCC1 | CN1CCN(C(=O)C2CCCO2)CC1 | CC(=O)N1C[C@H](O)C[C@@H]1C(=O)O |
| O=C(Nc1ccc(-n2cnnn2)cc1)C1CCC1 | O=C(CCc1ccccc1)Nc1ccc(O)cc1 | c1ccc(CCNc2nc3ccccc3[nH]2)cc1 | OCc1cn(-c2ccc(Cl)c(Cl)c2)nn1 |
| Cc1ccc(S(=O)(=O)N(C)CCC#N)cc1 | CN(C)c1ccncc1 | Oc1ccccc1CNc1nc2ccccc2[nH]1 | CNC(=O)C1CNCCO1 |
| c1ccc(-c2ccc(-n3cnnn3)cc2)cc1 | CCC(C)NC(=O)c1ccccc1N | CNC1CCN(c2ccccc2F)C1=O | c1cnc(NC2CCCCCC2)nc1 |
| O=C(Nc1cccnc1)c1cnn2cccnc12 | CC(C)c1ccc(CN2CCC(O)CC2)cc1 | COc1cccc(CN2CCC(C(N)=O)CC2)c1 | O=C(CCC1CCCCC1)N1CCOCC1 |
| CCNC(=O)c1ccc(N(C)C)c(F)c1 | Cc1ccc(Nc2ncnc3[nH]ncc23)cc1 | CNC1CCCN(c2cccnn2)C1 | O=C(Cc1ccc(Cl)cc1)Nc1ncccn1 |
| c1cn(CC2CCCCN2)nn1 | CCOc1ccc(CC(=O)O)cc1 | CNC1(CO)CCOCC1 | c1csc(-c2nnc[nH]2)n1 |
| O=C(Nc1cccc(F)c1)c1cnccn1 | C1CC(c2nc(C3CC3)no2)CN1 | CCNc1ccc(S(C)(=O)=O)cc1F | N#Cc1ccc(NC(=O)c2cccs2)cc1 |
| COc1ccc(Br)cc1CNC(C)=O | Fc1cccc(CNCc2ccco2)c1 | Cc1cc(Cn2nc(C)ccc2=O)on1 | O=C(c1ccc(Cl)cc1)N1CCSCC1 |
| NC(=O)c1cnn(CCO)c1 | O=C(Cc1ccc(F)cc1)Nc1ccc(F)cc1 | CNC(=O)c1cnc(C)s1 | COC(=O)c1ccc(NC(=O)C(F)(F)F)cc1 |
| CC1CCCCN1CCN | COC(=O)c1ccncc1 | CN(CCc1ccccn1)C(=O)c1ccccc1 | COCC(=O)NCc1ccc(C)cc1 |
| CS(=O)(=O)N1CCC(C(=O)NC2CC2)CC1 | CC(N)c1ccc(NC(=O)C2CC2)cc1 | Fc1cccc(CN2CCOCC2)c1 | CC1(C)CN(C(=O)CC(F)(F)F)CCO1 |
| Cc1ccc(CC(=O)NC[C@@H]2CCCO2)cc1 | CCC(CC)C(=O)Nc1nccs1 | CCOc1c(F)cccc1C(=O)N(C)C | N#Cc1ccc(N2CCCOCC2)cn1 |
| CC(=O)NCc1cn[nH]c1 | Nc1cc(C(F)(F)F)ccc1N1CCCCC1 | CN1CCN(C(=O)Nc2ccc(F)cc2)CC1 | CC1CCCC1NCCO |
| CCOc1ccc(C(=O)Nc2ccccc2)cc1 | CC(O)COc1ccc(F)c1 | Fc1ccc(Cn2cnc3ccccc32)cc1 | CC(=O)Nc1ccccc1O |
| NC(=O)c1c[nH]c(C2CC2)n1 | CC(C)C(=O)N1CCN(C(C)C)CC1 | CN1CCN(c2ncccc2F)CC1=O | CN(C)Cc1ccc(C(=O)O)o1 |
| CCOC(=O)C1CCN(c2cnccn2)CC1 | CC1CCN(C(=O)CCCc2ccccc2)CC1 | CC(=O)N1CCCC2(CCCCCC2)C1 | CC(=O)Nc1ccc(N2CCN(C)CC2)cc1 |
| N#CCC(=O)N1CCC(Cc2ccccc2)CC1 | c1ccc(SCCN2CCOCC2)cc1 | CC(N)c1nc(Cc2ccccc2)no1 | CC(=O)Nc1ccccc1CC(=O)O |
| CC(Oc1ccccc1F)C(=O)O | N#Cc1ccc(CNCc2ccc(F)cc2)cc1 | Cc1nc(C)n(CCC(=O)O)n1 | CC1CN(Cc2nnsc2Cl)C(C)CO1 |
| Cc1cc(N2CCNCC2)nc(C)n1 | Cc1cccc(Nc2ncnc3c2cnn3C)c1 | Cc1ccc(NC(=O)c2ccco2)cc1O | CC(Oc1ccc(C#N)cc1)C(N)=O |
| Cc1cccc(C(=O)N2CCC(C(N)=O)CC2)c1 | CCN1CCC(Nc2ccccc2F)CC1 | Cc1nc2ccccc2n1CCC(N)=O | COC(=O)CNC(=O)C1CCCCC1 |
| C[C@@H](CO)N(C)c1ncccc1F | O=C(Nc1nc2ccccc2[nH]1)C1CCCCC1 | CCOc1ccc(C(C)N)cc1 | FC(F)(F)c1ccc(CN2CCOCC2)cc1 |
| Cc1ncsc1C(=O)N1CCCNCC1 | CNC(=O)c1cccc(C(=O)O)n1 | CC1=NN(c2ccccc2)C(=O)C1 | CCOC(=O)c1ccoc1 |
| Cc1n[nH]cc1CNc1ccccc1F | Cc1cc(C(=O)NCC2CCOC2)on1 | Fc1cccc(CNCc2ccccc2)cc1 | CC(N)CN1CCOCC1 |
| c1ccc(CNC2CCNC2)cc1 | Cc1cccc(NC(=O)CN2CCOCC2)c1C | NC(=O)C(O)C1CCOCC1 | CCC(CC)C(=O)Nc1ccccc1O |
| Cc1ccc(CC(=O)Nc2nccs2)cc1 | CNC(=O)NCc1c(F)cccc1Cl | O=C(CC1CCCCC1)Nc1cccnc1 | COC(=O)C1CCN(CC(N)=O)CC1 |
| CCN(CC)c1cccc(O)c1 | c1csc(-c2nc(N3CCCCC3)no2)c1 | COc1ccc(C)cc1NC(=O)CN | NC(=O)C1CCC(F)(F)CC1 |
| CCc1nc(-c2ccc(Cl)cc2)no1 | Cc1csc(NCC2CCCO2)n1 | CN1CCN(C(=O)C2CCCN2)CC1 | CNC(=O)C1(Cc2ccccc2F)CC1 |
| O=C(COc1ccccc1)Nc1cccc(F)c1 | O=C(COc1ccccc1)Nc1ccccc1O | Cn1ccc(C(=O)NC[C@@H]2CCCO2)n1 | Cc1cc(C(=O)NC(C)C)nn1C |
| CCOc1ccc(NC(=O)NC(C)(C)C)cc1 | O=C(Nc1nncs1)c1ccccc1F | CC(=O)NCC1(c2ccccc2)CCOCC1 | COc1ccc(CC(=O)Nc2nncs2)cc1 |
| CN(Cc1cccc(F)c1)S(N)(=O)=O | CC1CN(C)CCC1Nc1cnn(C)c1 | Nc1ccc(S(=O)(=O)Nc2ccccn2)cc1 | CNS(=O)(=O)CC1CCCCC1 |
| CC(C)NC(=O)c1ccc2ccccc2n1 | COc1cc(NC(C)=O)c(OC)cc1Cl | CC1CCN(CCN)CC1 | COc1ccc(NCc2ccc(F)cc2)cc1 |
| COC(=O)c1ccc(NC(=O)C2CCCC2)cc1 | Cc1cc(C(=O)Nc2ccn(C)n2)on1 | CCCn1c(NC(=O)CC)nc2ccccc21 | Cc1ccc(CNC(=O)c2sccc2C)cc1 |
| N#CCCN1CCOCC1 | Cc1noc(C)c1CC(C)C(=O)N(C)C | CC(=O)Nc1cccc(NC(N)=O)c1 | COC(=O)Nc1nnc(CC2CCCC2)s1 |
| O[C@@H]1CCN(Cc2ccc(Cl)cc2)C1 | O=C(CN1CCCC1)Nc1ccc(F)cc1 | CC(C)C(=O)N1CCC(Cc2ccccc2)CC1 | CCOC(=O)c1sc(-c2cnccn2)nc1C |
| CCc1ncc(CNC)s1 | Cc1nn(C)c(C)c1NC(=O)CC#N | N#Cc1ccc(NC(=O)CCC2CCCC2)cc1 | CC(C)NC(=O)N(C)Cc1cnn(C)c1 |
| O=C(NCCc1ccncc1)c1ccccc1F | COCC(=O)Nc1ccccc1C(N)=O | CCc1ccc(C(=O)NCc2cnn(C)c2)s1 | CCC(NC)c1ccncc1 |
| COc1ccc(CCC2(C)NC(=O)NC2=O)cc1 | COCC(=O)NCc1nc2ccccc2[nH]1 | CN(C)C(=O)c1ccc(F)cc1Br | CC(C)c1nccn1CC(=O)O |
| O=c1[nH]cc(NCc2ccccc2)c(=O)[nH]1 | CC(=O)NCc1ccc(S(N)(=O)=O)cc1 | CC(=O)NCCN1CCCCCC1 | CC(=O)Nc1ccncc1C |
| Cc1ccc(NC(=O)c2cccc(F)c2)cc1O | CC(=O)Nc1ccc(NC(=O)CC(C)C)cc1 | Cc1ccc(CS(N)(=O)=O)cc1F | CNCCC1CCCCC1 |
| Cc1nccn1CC(C)O | O=c1[nH]cc(N2CCOCC2)c(=O)[nH]1 | Cc1ccccc1CNc1nnnn1C | OC1CN(C(c2ccccc2)c2ccccc2)C1 |
| CN1CCN(S(=O)(=O)N(C)C)CC1 | CC(NS(C)(=O)=O)c1ccccc1Cl | CC(=O)Nc1nnc(SC(C)C)s1 | CC(=O)N1CCOC(CO)C1 |
| COc1ccc(C(=O)C2CCNCC2)cc1 | CC1C(O)CCCN1Cc1ccccc1 | CC(=O)NCC1(c2ccccc2)C(=O)CC1 | Cc1cc(C2CCCN2S(C)(=O)=O)no1 |
| COCC(=O)Nc1nnc(C)s1 | CCc1nc(C2CCCN2)no1 | Cc1nn(Cc2c(C)noc2C)c(=O)s1 | CC(=O)N1CCC(c2c[nH]c3ccccc23)CC1 |
| CN(C)C(=O)c1cn[nH]c1 | NC(=O)c1ccc(NC(=O)c2cnccn2)cc1 | CCCC(=O)Nc1nnn(CC)n1 | CC1(S(=O)(=O)N2CCC=C(F)C2)CC1 |
| Cc1csc(CNC(=O)C2CCC2)n1 | CC(=O)Nc1ccc(-c2csc(N)n2)cc1 | O=C(Nc1ccccn1)c1cnccn1 | COCc1cc(C(N)=O)no1 |
| CC(=O)c1ccc(OCC(=O)O)cc1 | O=C(NCCN1CCCC1=O)Nc1ccccc1 | Cc1ccc(C)c(OC(=O)c2ccncc2)c1 | CCOC(=O)NCc1ccccc1 |

| | | | |
|---|---|---|---|
| CC1(C)CN(c2ncccc2F)CC1O | O=S1(=O)CCN(Cc2ccco2)CC1 | CCn1cc(NC(=O)C2CCC2)cn1 | Cn1cc(C(=O)O)c(-c2ccncc2)n1 |
| CC(C)c1ccc(C(=O)Nc2nncs2)cc1 | CSc1nsc(NC(C)=O)n1 | Cc1cc(N(C)C2CC(O)C2)nc(C)n1 | CN1CCC(Oc2cccc(F)c2)C1=O |
| CC1CN(C(=O)Cn2cccn2)CCO1 | CNc1ncc(C(F)(F)F)cc1Cl | O=C1CN(Cc2ccccc2)CC(=O)N1 | Cc1ccc(NC(=O)C2CCCO2)nc1 |
| CC(=O)NC1CCC(N)CC1 | O=C(COc1ccccc1)NC1CCCCC1 | CC(CS(N)(=O)=O)c1ccccc1 | Cc1cccc(C(=O)Nc2ccc(C)cc2O)c1 |
| CC(=O)NC(C)c1nnc2ccccn12 | CNc1ccccc1S(C)(=O)=O | CC(=O)Nc1ccc(Nc2ccccc2)cc1 | CC1CN(S(C)(=O)=O)CCC1N |
| CS(=O)(=O)N1CCCC(C(=O)O)C1 | CCn1ccc(CNC)n1 | O=C(NCc1ccc(F)cc1)Nc1ccccc1 | COc1ccc(NC2=NCCN2)cc1 |
| Cc1cccc(C(=O)NC(C)C2CC2)n1 | CNc1nccc(OC)n1 | CC(C)N1CCCC(CO)C1 | CC(=O)N1c2ccccc2CCC1C |
| NCc1ccccc1OC(F)(F)F | O=C(c1csnn1)N1CCOCC1 | COc1ccc2[nH]cc(CN(C)C)c2c1 | Fc1cccnc1NCC1CCOCC1 |
| c1ccc(Nc2nncs2)cc1 | Cc1ccc(CN(C)Cc2ccccc2)o1 | CCN(CC)C(=O)c1cc(C)no1 | CC(C)n1cnc(S(N)(=O)=O)c1 |
| O=C(O)CN1CCC(c2ccccc2)CC1 | COC(=O)Nc1ccc(Cc2ccncc2)cc1 | O=C(NC1CC1)c1cnn2cccnc12 | Nc1ncncc1-c1ccc(Cl)cc1 |
| OCC1CN(Cc2ccccc2)CCO1 | CC(=O)Nc1ccccc1OCc1ccccc1 | Fc1ccccc1CNC1CCOCC1 | CC1(C)CCCN1CCN |
| Cc1cc(F)ccc1CS(N)(=O)=O | CC(=O)NCCc1ccc(S(N)(=O)=O)cc1 | CN(C)C(=O)N1CCOCC12CCOC2 | COCCNC1CCN(C(C)=O)CC1 |
| Fc1ccc(NC2CCOCC2)nc1 | CC(=O)Nc1cc(C(N)=O)cc(C(N)=O)c1 | CC(=O)Nc1cccc2nonc12 | CN(CC1CCOC1)c1ncncc1Cl |
| Cc1nn(C)c(C)c1CC(N)=O | CC(=O)Nc1cnccc1C | CNCc1cccc(Cl)c1 | CN(C(=O)C1CCNC1)c1ccccc1 |
| Cc1ccn(Cc2csc(C)n2)c(=O)c1 | | | |
